# Interpretation of Phase Boundary Fluctuation Spectra in Biological Membranes with Nanoscale Organization

**DOI:** 10.1101/746800

**Authors:** S. S. Iyer, A. Negi, A. Srivastava

## Abstract

In this work, we use Support Vector Machine algorithm to detect simple and complex interfaces in atomistic and coarse-grained molecular simulation trajectories of phase separating lipid bilayer systems. We show that the power spectral density of the interfacial height fluctuations and in turn the line tension of the lipid bilayer systems depend on the order parameter used to identify the intrinsic interface. To highlight the effect of artificial smoothing of the interface on the fluctuation spectra and the ensuing line tension calculations, we perform a convolution of the boundaries identified at molecular resolution with a 2D Gaussian function of variance *ε*^2^ equal to the resolution limit, (1/2*πε*^2^)*exp*(*−|r|*^2^*/*2*ε*^2^). The convolution function is given by *h*⊗*g*, where *h* is the instantaneous height fluctuation and *g* is the Gaussian function. This is similar to the effect of point spread functions in experiments. We find that the region of fluctuation spectra that scales according to capillary wave theory formalism depends on the complexity of the interfacial geometry, which may not always be detected at experimental resolutions. We propose that the different q-regimes in the fluctuation spectra can be used to characterize mode dependent interfacial tensions to understand the interfaces beyond the linear line tension calculations. This could also be useful in interpretation of fluctuating boundaries in out-of-equilibrium *in-vivo* membrane systems that carry information about the nature of non-thermal (active) fluctuations in these systems.

## 1 Introduction

The study of lateral phase separation in the plane of the lipid monolayers and bilayer in terms of mobility of membrane anchored motifs and lipid conformations have come a long way since the early original work carried out 30-50 years ago [1–10]. Several recent experimental and computational studies have shown stark evidence of lateral phase separation in *in-vitro* model ternary [11–15] and quaternary lipid mixtures [16–18] and in *in-vivo* cell membranes [19–23]. The model ternary *in-vitro* systems generally consists of a saturated lipid, unsaturated lipid and cholesterol. Depending on the cholesterol content, ternary systems show two phase coexistence as “liquid-ordered” *L_o_* and “liquid-disordered” *L_d_* phases at higher cholesterol concentrations and “liquid-ordered gel” *L_β_* and “liquid-disordered” *L_d_* phases at lower cholesterol concentration. A three phase co-existence of *L_o_/L_d_/L_β_* has been observed at intermediate cholesterol concentrations [24]. Distinct phases differ in their chemical composition and physical properties.

Among the factors that dominate kinetics of phase separation and domain formation, line tension energy or the free energy cost per unit length of the interface, required to maintain the boundary is one of the major determinants of domain size [25,26], geometry and boundary roughness in lipid systems. The term was first introduced by Gibbs [27,28] as an analogue of surface tension acting at the domain interface. The energy at the domain boundary is given by the product of line tension and boundary length. Line tension can be tuned based on the chemical composition of the lipid mixture, which manifests as differential molecular interactions [16,29–32] and external factors such as temperature that effect distributions at the interface.

Similar to the thickness mismatch between lipid types in lipid-only systems, mismatch in the length of hydrophobic region of transmembrane protein and surrounding membrane in lipid-protein systems result in unfavourable interactions due to exposure of the hydrophobic region to aqueous environment. This excess energy per unit length is a major component of line tension. It is compensated by local membrane rearrangements of lipids around the protein [33,34]. Depending on the relative thickness, transmembrane proteins can create either local order, termed as orderphilic protein or local disorder, termed as orderphobic proteins [35]. The fluctuating interface between order and disorder phases generated by local rearrangements of lipids is characterized by its line tension. Minimization of line tension has been shown to be one of the key factors driving protein assembly in cell membrane [25].

Several experimental methods have been used to measure line tension at the boundary interface in lipid monolayer and bilayer systems. While AFM studies on lipid bilayer mixtures apply macroscopic classical nucleation theory to relate the nucleation rate of domains to line tension [36], line tension values measured using Flicker spectroscopy method employ capillary wave theory at microscopic length scales [37,38] and measurements using micropipette aspiration technique relate laplace pressure to line tension in the limit of the interfacial boundary energy being much larger compared to the bending energy [39,40]. Although the line tension values reported from these different methods are similar in magnitude for the same ternary lipid mixture studied [41], they are limited by resolution of the experimental methods applied. In this work we concentrate on the capillary wave theory formulation applied on domain boundary fluctuations to quantify line tension. Value of line tension obtained and important features of the boundary fluctuation spectra are highly dependent on the boundary identified.

Roughness of the boundary depends on the length scale at which it is measured. The molecular simulations trajectories do not have the resolution bottleneck and can be used to model and study the molecular-level roughness at the domain boundaries. In this work, we study boundary fluctuations in coarse grain lipid bilayer systems resulting from order-disorder interface generated due to protein insertion in the bilayer and due to phase separation in ternary lipid mixtures. These boundaries are identified using Support Vector Machine (SVM) algorithm and user-defined supervised classifiers. The highly automated machinery of SVM allows us a high-throughput rigorous quantification of the interface boundaries at molecular scale with different user-defined order parameters (classifiers) and allows us to look into the resulting fluctuations spectra, not just at the steady state but also while the domain formation is in progress.

This article is organized as follows. In Section II we describe the systems under investigation, discuss the methods and algorithms used to identify the boundaries and also discuss the capillary wave theory applied on the boundary fluctuations. In Section III, we show the results of application of SVM to identify boundaries in systems with simple and complex geometries and discuss the results with respect to the reported methods of intrinsic surface identification. Further we show the effect of using different order parameters in interface and their fluctuation spectra. Finally in this section we show the effect of resolution on the geometry of the identified boundary and its implications on the fluctuation spectrum and the line tension calculations. We conclude in Section IV by summarizing the major results in this work and put forth some speculations on how fluctuation spectra can be interpreted to understand non-equilibrium nature of systems.

## 2 Material and Methods

### 2.1 All-Atom (AA) and Coarse-Grained (CG) systems

We use five different systems, as listed in Table 1, for our study. The two All-Atom (AA) trajectories exhibiting fluid-phase coexistence [42,43] were made available by the Pittsburg Supercomputing Centre. There are three Coarse-Grained (CG) trajectories of lipid bilayers that are all simulated using Martini CG force-field parameters [44]. While the first CG system (DPPC/DUPC/CHOL ternary lipid mixture) is an example of fluid-fluid phase coexistence with a quasi-circular boundary [45], the second one is a transmembrane protein - lipid bilayer system (DPPC/TMP) with a circular interface geometry [35]. The third CG system is a transmembrane peptide - lipid bilayer (DAPC/DPPC/CHOL with 9 tLAT peptides) system with a linear interface geometry [46]. All CG trajectories were borrowed from the respective laboratories for implementation and analysis of the algorithms discussed in this paper. Simulation details about the systems can be obtained from the original papers where these trajectories were generated.

**Table 1:**
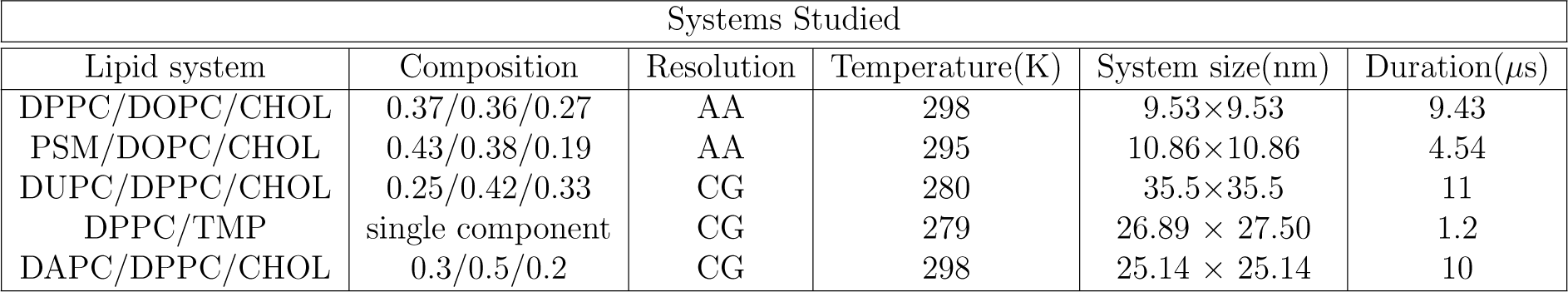
Phase separated systems for which boundary is identified

Analysis were carried out on the last 1.84 *µ*s of the trajectory of DPPC/DUPC/CHOL system after the *L_d_* lipids cluster into a quasi-circular lateral organization geometry. The last 2.3 *µ*s of the trajectory was split into 5 independent segments and fluctuations were calculated for each segment and averaged across the segments. For the DPPC-TMP system, fluctuations of the interface were calculated over 10 independent trajectories (600 *ns* each). For the DPPC/DAPC/CHOL systems, the interface fluctuations were calculated in the last 1 *µ*s of the trajectory for system with and without tLAT transmembrane peptides. For the AA systems, we have taken individual snapshots of the trajectory to detect domain boundaries and due to the highly rough and transient nature of the interface, fluctuation spectra analysis is not carried out on them.

### 2.2 Order parameters used as domain boundary classifiers

We apply membrane thickness, deuterium order parameter [2,4,47] and hexatic order parameter (*φ*_6_) [48] to characterize the extent of membrane orderliness for the simulation systems. Besides these commonly used markers, we also use the degree of non-affine displacements (*χ*^2^) [49] to distinguish between the different phases in the lipid bilayer systems. We recently showed that *χ*^2^ works as a high-fidelity marker between the liquid order (*L_o_*) and liquid disordered (*L_d_*) regions in the membrane system at molecular length scales [31,50,51]. *χ*^2^ was originally applied on granular material by Falk and Langer ([49]) and is given as:

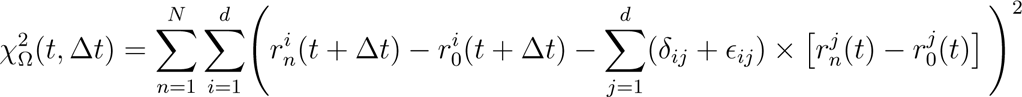

where the indices *i* and *j* run through the spatial coordinates for dimension *d* and *n* runs over the *N* lipids in the neighbourhood Ω, defined within a cutoff distance around the reference lipid *n* = 0. *δ_ij_* is the Kronecker delta function. *ϵ_ij_* is the strain associated with the maximum possible affine part of the deformation and thus, minimizes 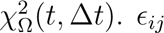 is defined as given in the original reference [49]. Δ*t* of 240 ps is used for AA systems and 8 *ns*, 4.6 *ns* and 60.0 *ns* for DPPC/TMP, DPPC/DUPC/CHOL and DPPC/DAPC/CHOL CG systems, respectively. Lipids in the *L_d_* phase have higher *χ*^2^ values than lipids in *L_o_* phase [31,50].

### 2.3 Detecting Boundaries using SVM

Identifying molecules at the phase boundaries and measuring the atomic scale motions to quantify the interfacial properties is a challenge in experimental and computational science. Line tension measured through fluctuations of the interface depends heavily on precise identification of the interface. In this work, we identify the boundary using SVM classifier method and ensure that the boundary thus identified is the true intrinsic interface. Thermal broadening of the intrinsic surface due to capillary waves results in the observed intrinsic width of the interface. Algorithms for the identification of intrinsic surfaces have come a long way from the original formulations where interfaces were assumed to be a step function [52]. These can be broadly categorized into methods that require the precise identity of interfacial molecules [53] and those which do not rely on identification of interfacial molecules and involve obtaining a smooth interface by spatial coarse graining [54]. In general, systems exhibiting fluid-phase coexistence do not have very clear interfacial region and detection of phase boundary becomes even more involved if the system has not yet reached a steady state, approaches critical point or is affected by external factors such as inclusions in the membrane matrix or external perturbations. In this work, we harness the highly automated SVM machinery to detect phase boundaries in bilayers exhibiting fluid-phase coexistence.

SVM [55,56] refers to a class of supervised learning algorithm that is used for data classification, regression analysis and outlier detection. With advent of readily available software tools, it is now extensively being used in the area of computational biology to predict protein structure [57], classify protein into functional family [58], recognize protein folds [59] and predict lipid interacting residues in membrane proteins [60] to name a few. For the sake of completion, we have described the algorithm in the SI. Broadly speaking, for a set of training data set with multiple classes, the purpose of SVM algorithm is to generate hyperplanes that maximally separates the data from one another. In this work, the data set is divided into two classes. Lipids with order parameter above a certain value are marked in one class while the ones below a certain cutoff values are marked in another class. SVM algorithms use set of mapping functions called kernel functions, which are particularly convenient to classify data that are linearly non-separable. In this work we use radial bias function (RBF, *K*(**x**, **x**’) = *exp*(*−γ||***x** *−* **x**’||^2^)) and linear kernel function (*K*(**x**, **x**’) = **x**.**x**’) upon appropriate optimization of the parameters. These are implemented on SciPy tools [61] and scikit-learn contains the machine learning library [62] in python. We have put our code with example input files on github (Provide link here)

### 2.4 Fluctuation Spectra and Capillary Wave Theory

Capillary Wave Theory (CWT), which draws correlations between capillary wave fluctuations and interfacial tension [52,63,64], presents a method to calculate interfacial tension from the power spectral density of interface height fluctuations without the knowledge of forces between individual molecules. CWT assumes that the interface is acted upon by thermal waves causing small amplitude fluctuations and gives a direct connection between the amplitude of the fluctuations of the intrinsic surface and line tension associated with the boundary [52]. The small amplitude fluctuations contribute to quadratic free energy function for these fluctuations. Thus the mean square amplitude of Fourier components of the capillary waves follow a Gaussian distribution and each modes contribute *k_B_T*/2 energy according to equipartition theory, within the equilibrium approximation. Mode dependent interfacial line tension is given by [65]:

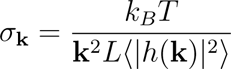

where *k_B_* is Boltzmann constant, *T* is temperature, *L* is perimeter of the boundary, 〈*|h*(*k*)*|*〉 is the power spectral density of height fluctuations of the interface and k is 2*πn/L*. This is commonly used in fluorescence microscopy of liposomal vesicles exhibiting domain separation [37]. While the above formulation is applied to vesicular domains by getting the power spectra of the height fluctuation (radial deviation) about mean perimeter (mapped to a planar boundary of equal length), another variant of the above fluctuation spectra formula commonly used in Flicker Spectroscopy to obtain line tension [38] is applied for circular / quasi-circular interfaces. In the former case, with planar interface boundary representation, the Hamiltonian of the interface is written in cartesian co-ordinates and Fourier transformation of the height fluctuations is given in a interval between 0 and L. For systems with circular / quasi-circular interface, the periodicity is 0 to 2*π*. The Hamiltonian of the interface is written as a function of arc length in polar coordinate system. Fourier series of the radius (height) fluctuations is written with a periodicity of 2*π*. Brown and co-workers have beautifully laid out the theoretical underpinnings of the domain fluctuations to membrane dynamics, viscosity and molecular orientations and dynamics of lipids [66,67]. We show the equivalence of the two formulations in SI and have applied both methods for line tension calculations.

Once the boundaries are appropriately detected at all snapshots, we calculate the fluctuation spectra of the in-plane boundary fluctuations and study its properties. To subtract for the effect of translation on the identified boundary, we rescale the boundary co-ordinates with respect to the centre of the boundary for every configuration analyzed from the molecular dynamics trajectories. Further, to obtain a smoother boundary, we interpolate across the boundary points using B-spline algorithm. Once the set of boundaries are collected, the overhangs are discarded in the fluctuation analysis. For circular boundaries, instantaneous height fluctuations are given as: *h*(*θ*) = *H*(*θ*) − 〈*H*(*θ*)〉 where *h*(*θ*) is the instantaneous radial deviation from mean radius 〈*H*(*θ*)〉 with similar corrections made for linear boundaries. To obtain 〈*|h*(**k**)*|*^2^〉, discrete Fourier transform of *h*(*θ*), obtained over 460 *ns* (100 frames, Δ t = 4.6 *ns* CG timescales) for DPPC/DUPC/CHOL system and 600 ns for DPPC/TM system, was performed. Fluctuations were also calculated from other segments of same time frames for both the systems to ensure proper statistical averaging. The boundaries obtained from each segment and their average is shown in Fig. S1 (chemical identity as classifier) and Fig. S2 (tail order parameter as classifier). The need for long trajectories arises because of the long timescale dynamics of the larger wavelengths (small **k**) modes. These modes are difficult to capture with statistical accuracy. 〈*|h*(**k**)*|*^2^〉 value was again averaged over the independent segments of trajectories for statistical accuracy. Averaging across frames in a trajectory and across independent sets of trajectories is performed to obtain the of power spectrum density (PSD). PSD thus obtained is a Fourier transform of autocorrelation of amplitude of height fluctuations.

## 3 Results and Discussion

### 3.1 Uncertainty Quantification of Interfaces using SVM

Precise quantification of continuous interface boundary between coexisting fluid phases is an involved exercise at molecular scales since the interface is under thermal fluctuations that constantly affects the local and global density profiles around the interface. Detecting phase boundaries in a non-equilibrium system or system that is evolving and on the way to equilibrium is even more challenging due to the highly dynamics nature of the interfaces. In the past, several schemes have been attempted to faithfully capture the continuous interface boundaries in lipid bilayer systems with coexisting *L_o_* and *L_d_* phases. Carla Rosetti and co-workers [30] obtained phase boundaries by using the intrinsic density profile method [53] after eliminating the thermal fluctuations of the interface. David Chandler and co-workers [35] used Gaussian density field approach [54], where they perform a convolution with a Gaussian function and smoothen out the interface profile. Voronoi tessellation with a cut-off on tail order parameter [68] and local composition [16] have also been applied to identify lipids in *L_o_* and *L_d_* phases and their the phase boundaries. Recently, John Straub and co-workers used an interface detection algorithm, similar to that used by Carla Rosetti and co-workers, that uses the information about the chemical identity of a given lipids nearest neighbours to develop the interface profile [41].

We leverage the highly automated machinery of SVM method to identify domain boundaries that allows us to quantify the inherent uncertainty in detection of interfaces. Given labelled training data set (supervised learning), SVM outputs an optimal hyperplane that maximally separates the data with different labels. For example, if the phase boundaries have to be detected using lipid type, we label saturated lipids with +1 label and unsaturated lipids with *−*1 label and use the SVM algorithm to identify boundaries. Fig. 1 shows the boundaries detected for a single snap shot of the AA DPPC/DOPC/CHOL system, PSM/DOPC/CHOL system and CG DPPC/DAPC/CHOL system with and without the tLAT peptide. While DPPC/DOPC/CHOL system has a closed-loop boundary, the PSM/DOPC/CHOL system has a linear boundary. For systems with closed boundaries, SVM applies a Radial Bias Function (RBF) kernel given as *exp*(*−γ||x − x*’||^2^) on the pre-classified data, to find the optimally separating hyperplane. For linear interface, linear kernel suffices to distinguish the *L_o_* and *L_d_* phases and obtain the boundary. The *γ* parameter determines the spread of the Gaussian and hence the range of influence of the support vectors. Fig. 1 (1A-1B and 2A-2B) show two boundary profiles per AA system identified using different values of *γ*. Depending on the *γ* parameter value (see Table 2), we show two different boundary profiles per CG system and further discuss the sensitivity of boundary profile to SVM parameters shortly. Movie movS1a and movS1b in SI show the boundary detected using chemical identity for DPPC/DAPC/CHOL systems with planar interface [46]. As evident from the movies for the DPPC/DAPC/CHOL systems with planar interface, the boundaries are captured faithfully by the algorithm despite the noticeable interface fluctuations. The high fidelity of interface detection is also aided by the fact that the lipid miscibility across the interface is nominal, which need not be the case for all systems. For example, in movie file movS2a in SI, we show the near-circular domain formation kinetics process for a large DPPC/DUPC/CHOL bilayer system that starts from a random organization [45]. Movie file movS2b in SI shows the system with SVM-detected boundary fluctuations when the steady state is reached. In Fig. S1, all boundaries for last 900 snapshots are plotted for the DPPC/DUPC/CHOL bilayer with quasi-circular domain and in Fig. 2(A) we plot the boundaries for 100 snapshots (at 4.6 *ns* intervals) with the average boundary shown in black. One of the advantages of this highly automated method is the ability to track domain boundaries for evolving system since the method does not assume an equilibrated standing wave form at the interface. In Fig. 2(B), we show how the boundary length changes as a function of trajectory evolution. As expected, we do see that boundary length decreases with time (and becomes less jagged with time). However, the boundary length do not attain a constant value within the simulation time of 11 *µ*s of CG simulations. Convergence of the trajectory to its equilibrium configuration is not assured despite a visibly well-formed domain. We also discuss the repercussions of this with respect to the line tension calculations below.

**Figure 1:**
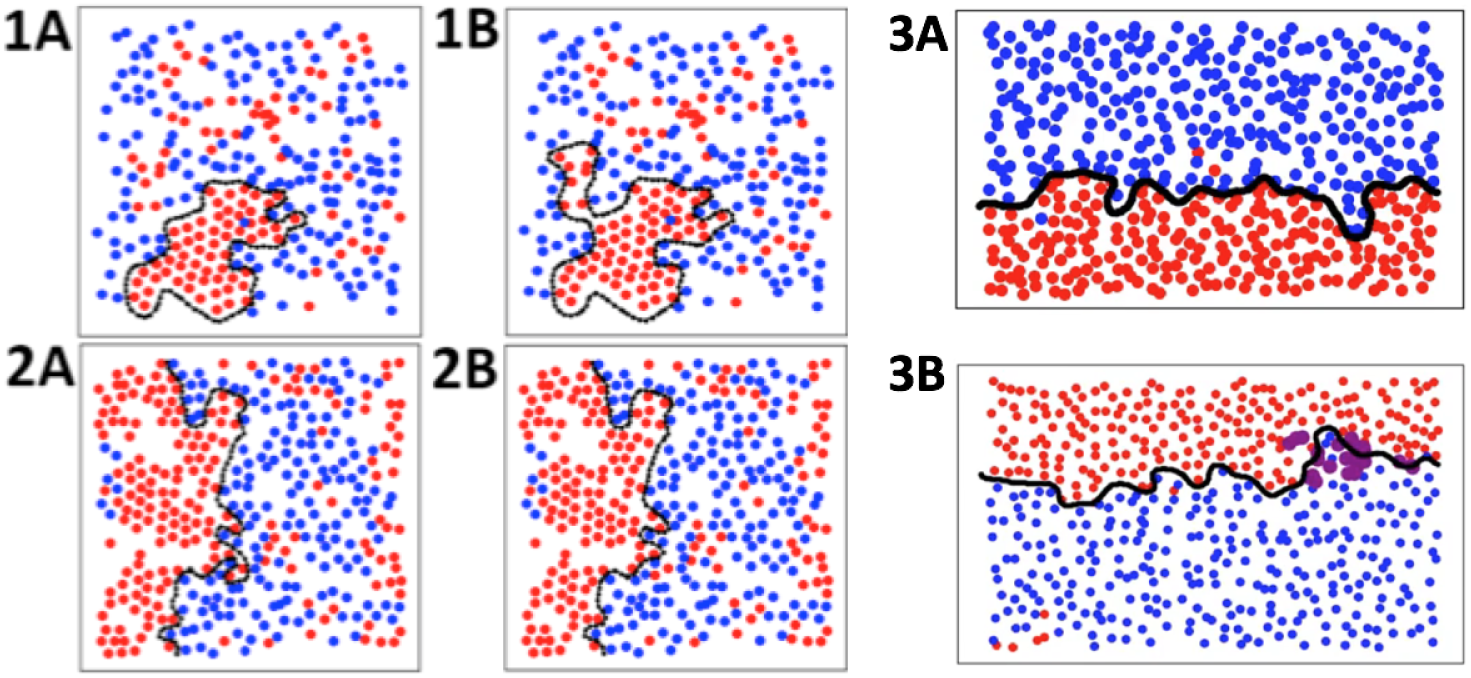
Boundaries determined for AA systems (1) DPPC/DOPC/CHOL and (2) PSM/DOPC/CHOL, and Martini CG systems (3) DPPC/DAPC/CHOL and (4) DPPC/DAPC/CHOL with tLAT peptide using SVM. (1A and 2A) corresponds to boundary determined with gamma value of 0.1 and (1B and 2B) corresponds to boundary determined with gamma value of 0.05. Saturated lipid sites are shown in red, unsaturated lipid sites in blue and boundary identified in black. The tLAT peptide localizing at the Lo/Ld interface in DPPC/DAPC/CHOL systems is shown in purple (3B).

**Figure 2:**
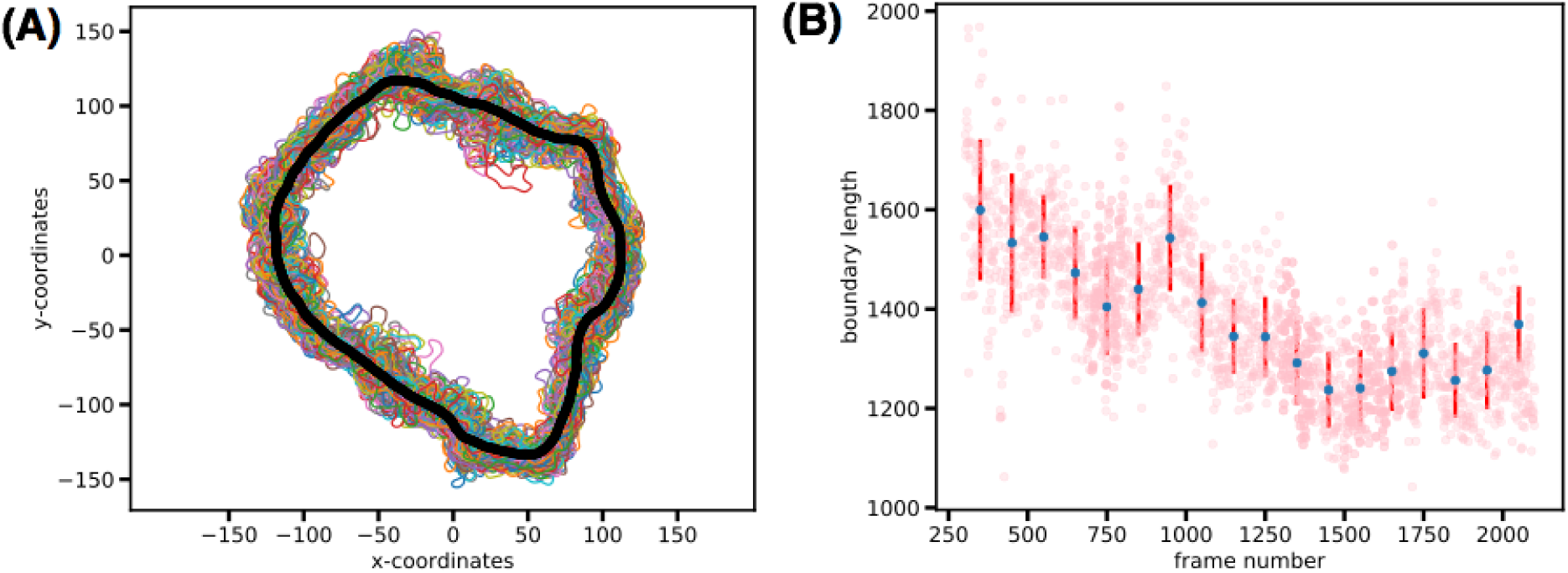
(A) Boundaries determined for CG DPPC/DUPC/CHOL system with quasi-circular interface using SVM. Chemical identity of the lipids is used to distinguish between the Lo and Ld phases. Boundaries for the last 100 frames are plotted and their average is shown in black. (B) Evolution of boundary length shown as a function of frame intervals. Boundary length measured at each frame is shown in pink points. As the simulation advances the boundary length is minimized. Blue dots shown the average boundary length for the frame range and the red lines determine the standard deviation.

**Table 2:** Properties of boundaries identified for phase separated DUPC/DPPC/CHOL and DPPC/TMP systems using different *γ* values: boundary width at maximum probability and variance in the probability distribution of boundary widths.

| Systems | $\gamma$ | $\sigma$ | w at $P_{max}$ |
| --- | --- | --- | --- |
| DUPC/DPPC/CHOL | 0.05 | 7.88 | 0.89 |
| DUPC/DPPC/CHOL | 0.1 | 9.35 | 7.73 |
| DPPC/TMP | 0.05 | 3.31 | 0.23 |
| DPPC/TMP | 0.1 | 4.61 | 1.10 |

Optimizing the value of gamma parameter in the SVM method is comparable to tuning the bin width resolution in the originally proposed intrinsic surface identification method by M. Jorge and M. N D.S. Cordeiro [53]. The value of gamma hence determines the lower wavelength cutoff of the capillary wave induced fluctuations on the intrinsic surface [69] and must be set to length scales of particle diameter. Small value of gamma implies higher variance of the Gaussian function and results in a smooth, less corrugated hyperplane or decision boundary. Large value of gamma implies a smaller range over which support vector *i* can influence the class of support vector *j* i.e, only if they are in close proximity. While setting gamma parameter to a very low value would imply incorporating particles that are otherwise part of the bulk phase rather than the particles that belong to the interface to mark the domain boundary. On the other hand, a high value of gamma could result in potential overfitting of the boundary. Hence different values of gamma will affect the shape of the boundary determined. This can be clearly seen in Fig. 1(1A,1B) and 1(2A,2B) which show different interfaces identified by SVM for the same systems with *γ* values of 0.1 and 0.05, respectively.

In order to identify which of the gamma values give the intrinsic boundary profile, we plot the probability distribution of widths of the fluctuating boundaries identified as shown in Fig. 3. The top panel represent the DPPC/DUPC/CHOL system with interface identified using chemical identity, while the bottom panel represent the DPPC/TMP system where a physical classifier (*χ*^2^) is used label boundaries. We discuss the physical classifiers in the next section. The width at maximum probability and variance of the probability distributions fit to Gaussians corresponding to Fig. 3 are given in Table 2. For rest of the analysis we use *γ* value of 0.05 which consistently shows lower variance of probability distribution and has a maximum for probability distribution occurring at w *≈* 0. This value of w signifies that the profile of interface obtained is *intrinsic* [30].

**Figure 3:**
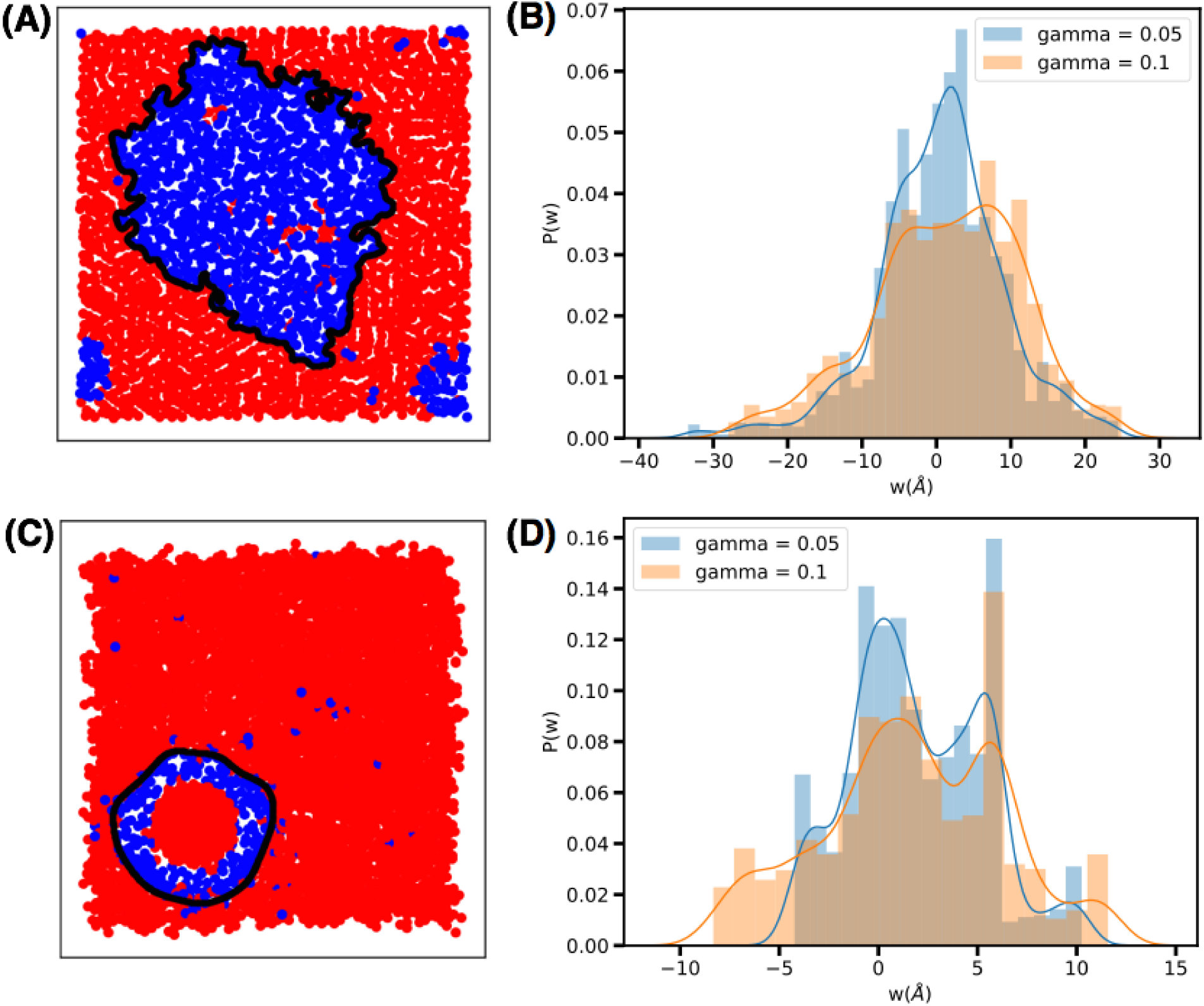
(A) and (C) show the interface identified for Martini CG DPPC/DUPC/CHOL and DPPC/TMP systems using SVM. (C) and (D) show the probability distribution of widths of boundary identified with *γ* of 0.05 and 0.1 for the corresponding systems. *γ* value of 0.05 shows maximum probability of width closer to 0 and lesser variance in the width distributions, indicating that it can be used to identify intrinsic surface.

### 3.2 Choice of Order Parameter and Boundary Profiles

SVM is not restricted to using chemical identity as an order parameter to identify *L_o_* and *L_d_* phases. We have also used physical order parameters such as deuterium order parameter (*S_cc_*), non-affine displacement field (*χ*^2^), hexatic order parameter (*ϕ*_6_) and membrane thickness (*d*) as classifiers to detect phase boundaries with varying degree of sensitivity. For the DPPC/DUPC/CHOL system with closed-loop domain formation, Fig. S2 shows the domain boundaries for 900 snapshots with *S_cc_* as the classifier. The corresponding trajectory of domain formation is shown as movie file (movS3A in SI) where each lipid is colour coded as per its *S_cc_* value. For boundary identification, we choose a cut-off of *S_cc_* values, and label the lipids as *L_d_* for *S_cc_* value less than 0.5 and as *L_o_* for *S_cc_* value greater than or equal to 0.5 as shown in movie file movS3B in SI. The corresponding boundary at every snapshot is shown in movie movS3C. Same kind of evolution is also shown for *χ*^2^ (movie file movS4(a-c)) where we have used a cutoff of 2000 to classify *L_o_* and *L_d_* lipids.

The domain boundary and its fluctuation thus obtained with each of these order parameters is different. Fig. 4(A) and Fig. 4(C) shows the converged phase separated domain with each lipid color coded as per its *S_CC_* value and *χ*^2^ value, respectively. The difference in the average boundary for *S_cc_* cutoff value of 0.4 and 0.5 is shown in Fig. 4(B) and *χ*^2^ values of 2000 and 2400 is shown in Fig. 4(D). In Fig. 5, we plot the full fluctuation spectra of the interface. The difference in boundary obtained using different order parameters is reflected in the full fluctuation spectra shown in Fig. 5(A). The region between **k** value of 1 and 3 *nm*^−1^ scales as 1/*k*^2^, in accordance to CWT. Line tension values obtained by fitting these regions (as shown in Figure 5B) are 1.78 pN, 1.70 pN and 2.1 pN for fluctuation spectra of boundary obtained with chemical identity, *χ*^2^ (cutoff values of 2000) and *S_cc_* (cutoff value of 0.5) to distinguish *L_o_* and *L_d_* phases respectively. For *χ*^2^ cutoff value of 2400 and *S_CC_* cutoff value of 0.4, we find the corresponding line tensions values to be 1.07 pN and 1.25 pN, respectively. Fluctuations of square amplitude scaled by the boundary length lesser than 10^−3^*nm*^3^ in **k** region larger than 10 *nm*^−1^ contribute very minimally to the fluctuation spectra and can be ignored. Smaller **k** regions, corresponding to modes that are more difficult to equilibrate, scale as 1/*k^α^*, where *α* is a constant smaller than 2. These modes do not obey fluctuation-dissipation according to CWT and we discuss this further below.

**Figure 4:**
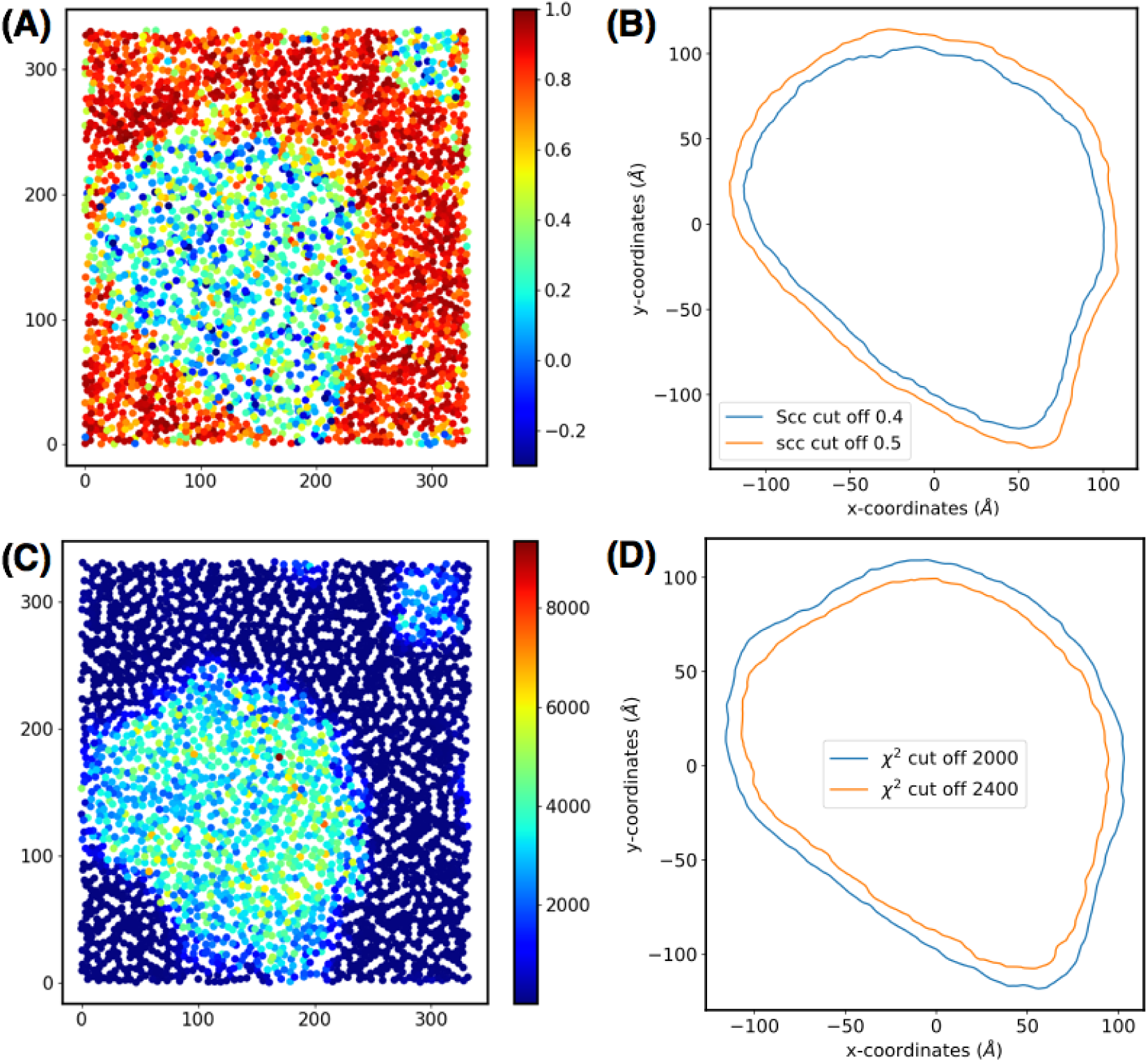
DPPC/DUPC/CHOL system mid carbon sites coloured according to (A) *S_cd_* values (C) *χ*^2^ values. system obtained using chemical identity, *χ*^2^ and *S_cd_* values. (B) and (D) show average boundary determined by a SVM using a cut-off of 0.4 and 0.5 for *S_cd_* values and 2000 and 2400 for *χ*^2^ values.

**Figure 5:**
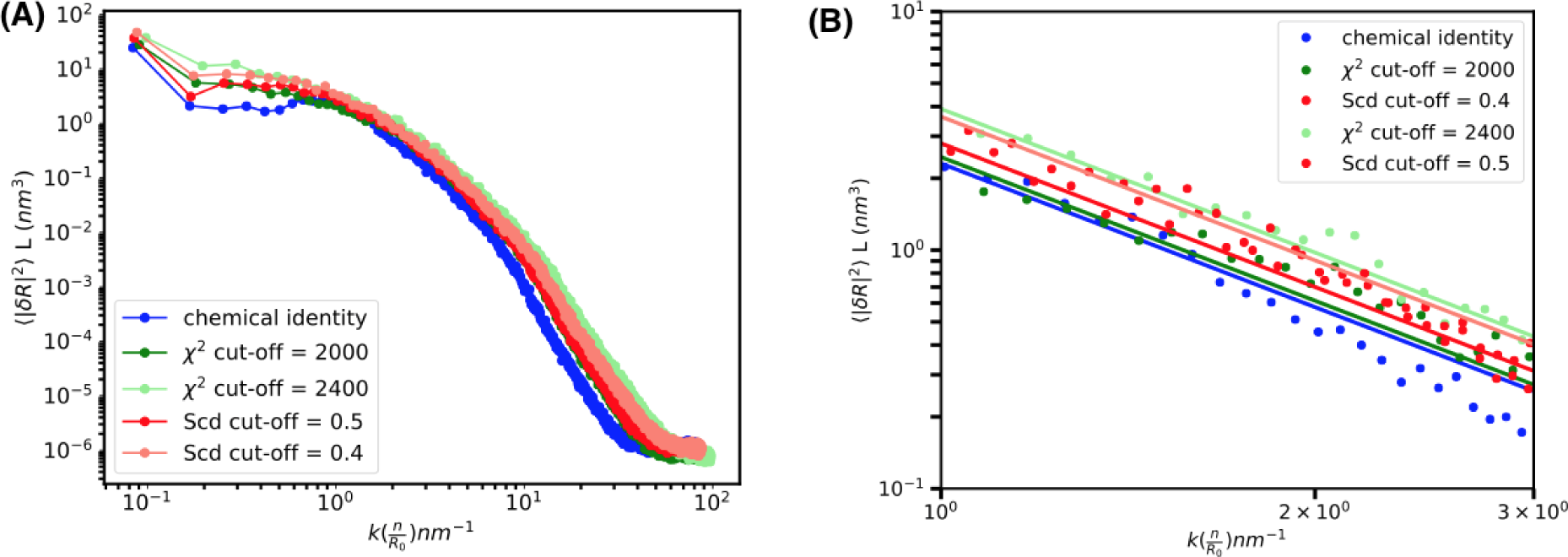
(A) Full fluctuation spectra for domain boundary for DPPC/DUPC/CHOL system obtained using chemical identity, *χ*^2^ and *S_cc_* values. (B) Fitting k region of 1 to 3 *nm*^−1^ in spectra obtained using chemical identity, *χ*^2^ and *S_cd_* values to capillary wave theory to obtain line tension values. Line tension values obtained using chemical identity is 1.78 pN, using *S_cc_* cut-off of 0.4 and 0.5 is 1.25 pN and 2.1 pN respectively, and using *χ*^2^ cut-off of 2000 and 2400 is 1.70 pN and 1.07 pN respectively.

Our analysis of the DAPC/DPPC/CHOL system (with and without tLAT peptide) shows that the line tension of the system decreases with the addition of tLAT peptides, which corroborates the free energy studies carried out on the system [46]. Figure 6 shows the fluctuation spectra and its fit for boundary marked used chemical identity and *S_CC_* to distinguish between the *L_o_* and *L_d_* phases. While the spectra obtained using fluctuations of the boundary identified using chemical identity and tail order parameter show a decrease in line tension upon insertion of the tLAT peptide, the magnitude of decrease is different in both these cases. Line tension obtained by fitting the **k** region between 1 to 3 *nm*^−1^ decreases from 4.50 pN to 2.12 pN and from 3.22 pN to 1.11 pN on localization of tLAT peptide at the interface for fluctuations of boundary obtained using chemical identity and tail order parameter, respectively. The above examples clearly show the effect of choice of order parameter and the cutoff used in the evaluation of line tension in the system. We also find (Fig. 7) that for molecular scale organization and for systems not having a very well-separated phase boundaries, thickness and hexatic order parameter are not the most robust classifiers to detect boundaries. In Fig. 7(A) and Fig. 7(C), we plot the planar distribution of local membrane thickness values and lipid-wise *ϕ*_6_ values, respectively for the DUPC/DPPC/CHOL system discussed above. Since the thickness calculation is not lipid wise and the thickness is calculated by interpolation of the grids on xy plane according to the GridMAT-MD algorithm [70], the local lipid-level order and disorder are highly homogenized. Fig. 7(B) and Fig. 7(D) show the *L_o_* and *L_d_* phases identified based on a cut-off on thickness and *ϕ*_6_ respectively. While *ϕ*_6_ is a good marker to distinguish between gel-fluid co-existing phases, we find that it is a poor marker to distinguish liquid-liquid co-existing phases [50]. The boundary marked in black in Fig. 7(D) is very rough. In another gel-liquid transition, Chandler and coworkers [35] perform a Gaussian convolution on the interface identified using *φ*_6_. The variance of the Gaussian used is in the order of molecular diameter of the lipid species. The corresponding trajectory evolution for thickness and *phi*_6_ are shown in SI movie files movS7A and movS7B, respectively. The *phi*_6_ works well for the above systems since the transition is gel-liquid but the Gaussian smoothening has some implications on the eventual spectra and line tension calculations, which we discuss in the next section.

**Figure 6:**
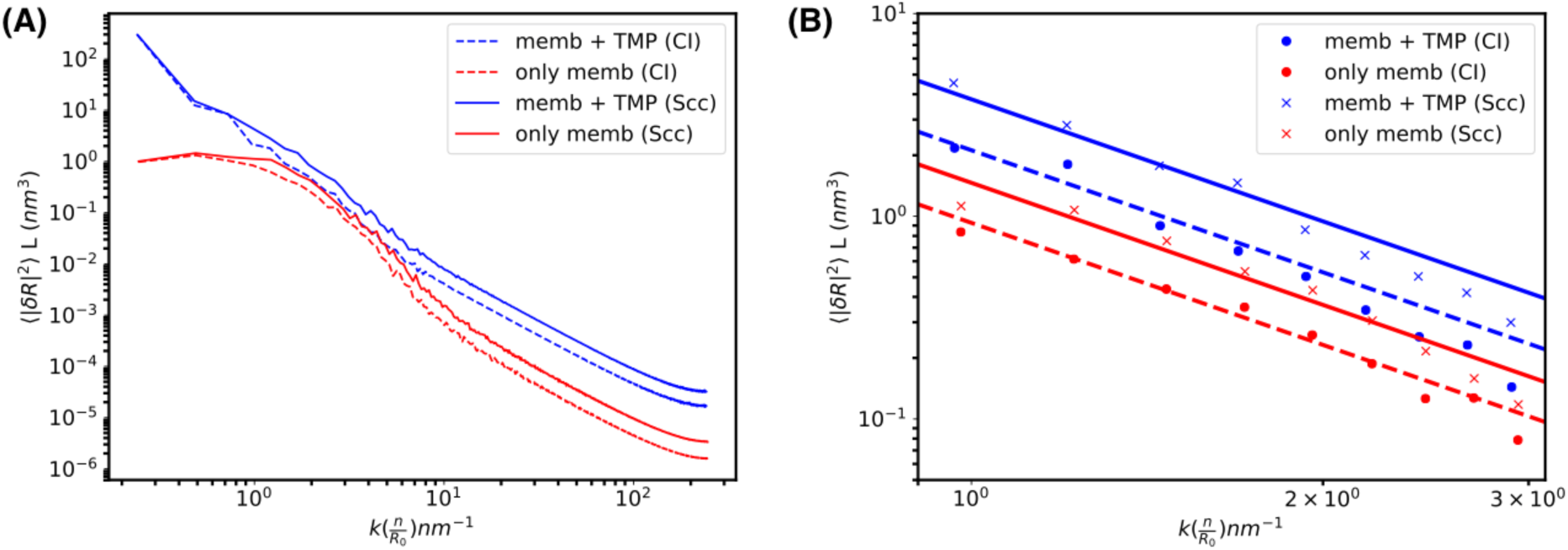
(A) Full fluctuation spectra for domain boundary for DAPC/DPPC/CHOL system obtained using chemical identity and *S_cc_* values. Red lines indicate membrane only system and blue line for the system with tLAT trans-membrane peptide. Solid lines represent fluctuations of boundary obtained using *S_cc_* order parameter and broken lines represent fluctuations of boundary obtained using chemical identity. (B and D) Fitting k region of 1 to 3 *nm*^−1^ in spectra obtained using chemical identity and *S_cd_* values to capillary wave theory to obtain line tension values. Line tension decreases from 4.50 pN to 2.12 pN and from 3.22 pN to 1.11 pN on localization of tLAT peptide at the interface for fluctuations of boundary obtained using chemical identity and tail order parameter, respectively.

**Figure 7:**
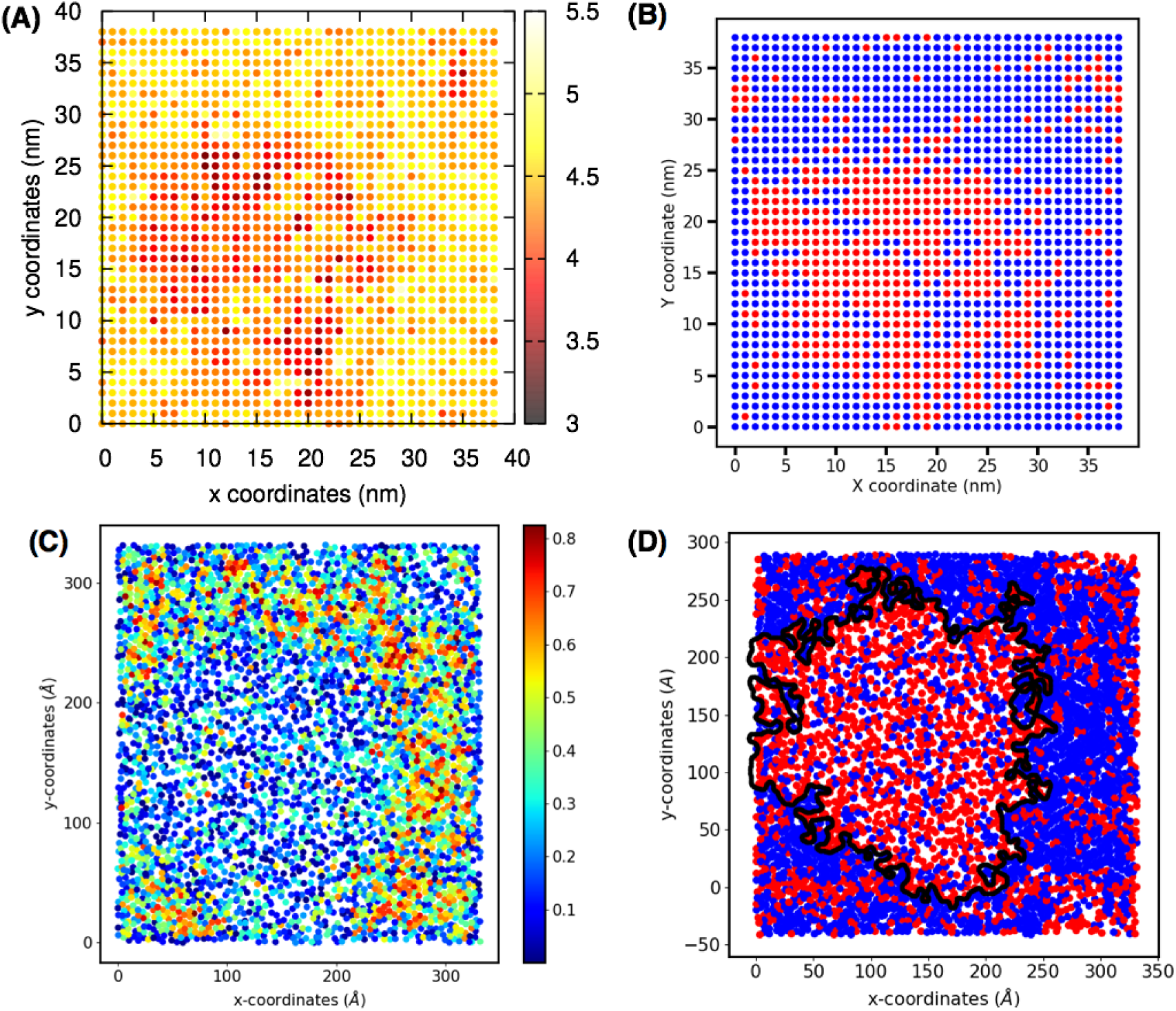
DPPC/DUPC/CHOL system grid points according to (A) thickness values and mid carbon sites coloured (C) *ϕ*_6_ values. (B) Lo and Ld phases classified based on a cut-off of thickness value. Grid points coloured in red indicate Ld phase and grid points in blue indicate Lo phase. (D) Lo and Ld phases classified based on a cut-off of *ϕ*_6_ values. Boundary identified using a cut-off of 0.12 of *ϕ*_6_ value, marked in black, gives a very rough interface.

### 3.3 Interpreting the complete boundary fluctuation spectra

Boundaries and their fluctuations obtained from molecular dynamics trajectories have resolutions up to molecular level details. Results from calculations performed on molecular dynamics trajectories need to be interpreted in terms of experiments such asflicker spectroscopy, which have limited optical resolution that can go as low as 86 *nm* [71–74]. To study the effect of artificial smoothing of the boundary observed due to resolution effects, we perform a convolution of the boundary identified at molecular resolution with a 2D Gaussian function of variance *ε*^2^ equal to the resolution limit, (1/2*πε*^2^)*exp*(*−|r|*^2^/2*ε*^2^), similar to the effect of point spread functions in experiments. The convolution function is given by *h* ⊗ *g*, where *h* is the instantaneous height fluctuation and *g* is the Gaussian function. We apply this prescription on (i) the DPPC/DUPC/CHOL system with quasi-circular boundary that is heavily discussed above, and also on (ii) the DPPC/TMP system [35] system. Boundary for the DPPC/TMP system is marked based on a cut-off on *χ*^2^ values. In this system chemical identity cannot be used to distinguish order and disorder phases (Fig. S3). Fig. S3(A) shows a snapshot with lipids sites with *χ*^2^ greater than 2000 labelled as *L_d_* in blue and lipids/TMP sites with *χ*^2^ lesser than or equal to 2000 labelled as *L_o_* in red color. The SVM determined boundary is marked in black. Fig. S3(B) shows the boundaries detected for 300 snapshots.The average boundary, shown in black, is almost a perfect circle for this system. The corresponding trajectory of boundary fluctuations is shown in movS6 in SI. Movie movS9 shows boundary marked and its fluctuation.

Fig. 8(A) and Fig. 8(C) show the effect of convolution on the boundary of DPPC/ DUPC/ CHOL and DPPC/TMP system, respectively. The effect of convolution is shown with a Gaussian function of variance 10 nm and 86 nm for both systems. As seen from the figure, the effect of smoothing with Gaussian of variance 86 nm is more apparent on the boundary of DPPC/DUPC/CHOL which has a highly rough interface at molecular-scales. The effect of Gaussian smoothening on the boundary profile results in an apparent quasi-circular domain shape in experimental images. While the roughness of the boundary is preserved on convolution with Gaussian of variance of 10 nm corresponding to an experiment with finer resolution, the corrugations in domain boundary is completely lost on performing a convolution with Gaussian of variance 86nm as seen in Fig. 8(A). The effect of smoothing of domain boundary is also easily captured by the fluctuation spectra shown in Fig. 8B for the DPPC/DUPC/CHOL system and Fig. 8(D) for the DPPC/TMP systems. Similar to the spectrum of DPPC/DUPC/CHOL system discussed above, fluctuations of square amplitude scaled by the boundary length lesser than 10^−3^*nm*^3^ in **k** region larger than 1 *nm*^−1^ contribute very minimally to the fluctuation spectra in DPPC/TMP system and can be ignored. The line tension values calculated for the spectra obtained by the rough boundary fluctuations and smoothened boundary fluctuations for the DPPC/DUPC/CHOL are 1.78 pN (*ϵ* = 10*nm*) and 31.08 pN (*ϵ* = 86*nm*). This is a significant increase (almost 20 times) and indictaes that smoothing of interfaces (particularly those with not well-separated interfaces) may lead to drastic over estimation of line tension values. For the DPPC/TMP systems, which has an almost circular average boundary (see Fig. S3B), the line tension values are found to be 11.84 pN (*ϵ* = 10*nm*) and 20.87 pN (*ϵ* = 86*nm*). The reported value of 11.84 pN for the fluctuations of intrinsic boundary is comparable to the value of 11.5 pN obtained by Shachi Katira et al [35] for the same system.The **k** region that is fit to obtain the line tension values for both these systems is shown in the inset of Figure 8(C) and 8(D) for DPPC/DUPC/CHOL system and DPPC/TMP, respectively.

**Figure 8:**
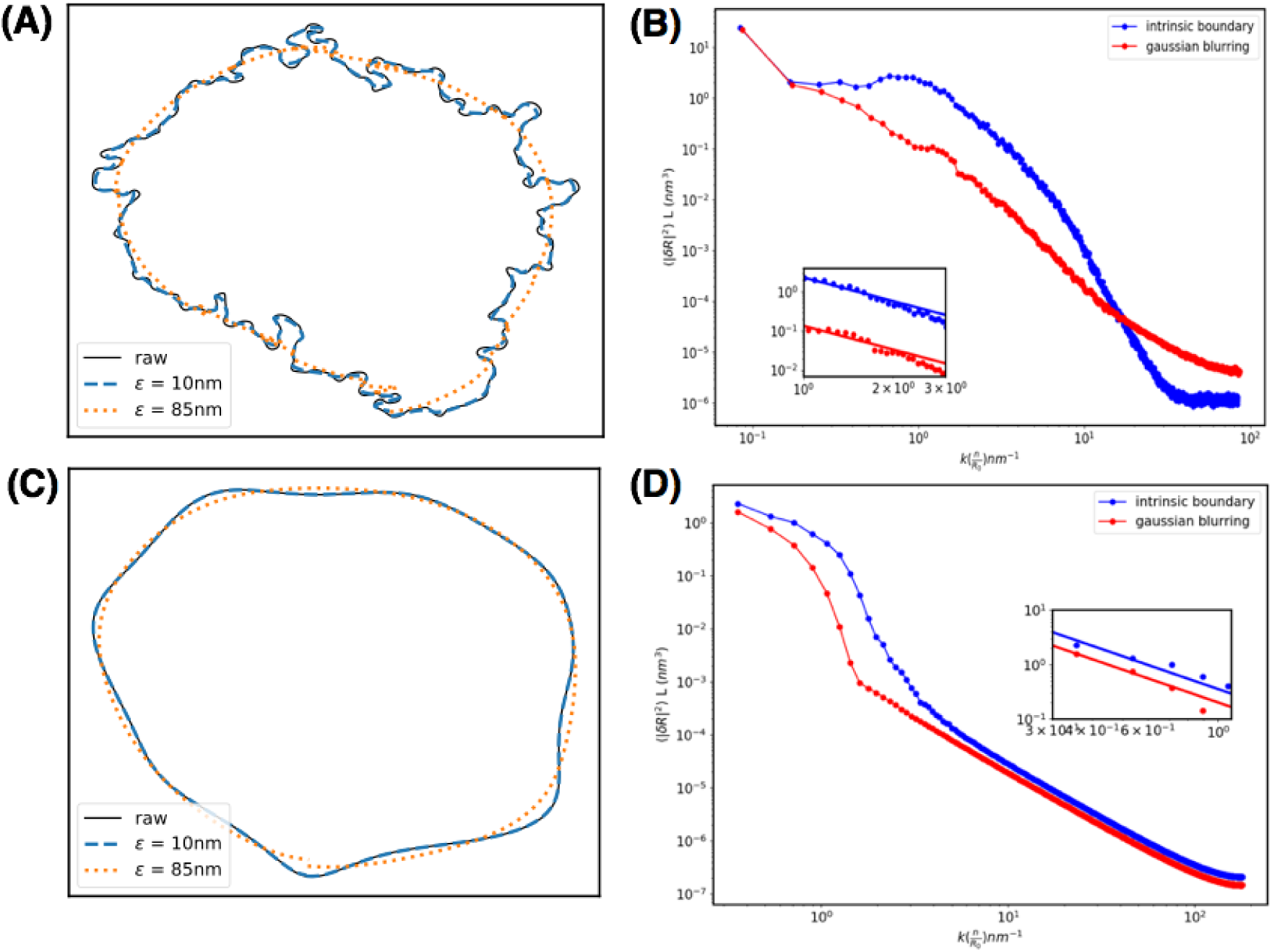
(A) and (C) Boundary of DUPC/DPPC/CHOL and DPPC/TMP system smoothened by convolution of a Gaussian function of variance 10 nm and 86 nm respectively. Effect of Gaussian smoothing on fluctuation spectra of (B) DUPC/DPPC/CHOL and (D) DPPC/TMP systems. Inset shows the k region fit to extract line tension values.

In Fig. S4, we show the spectrum obtained by an equivalent formalism (Formalism 2) [38,66,67] for the DPPC/TMP system that retains the circular reference geometry of the interface for power spectra calculations. The same first five **k** values are fit to CWT as shown in inset of Fig. 8(D). The line tension obtained is 10.01 pN as opposed to 11.84 pN by fitting the fluctuation spectrum to c/**k*^2^*** in the previously used formalism (Formalism 1) [37]. While formalism 1 does not impose strictly that the interface needs to be circular, formalism 2 assumes a perfectly circular interface. The difference in the line tension values obtained using the two formalisms could be due to the slight deviations of individual boundaries from non-circular geometries as shown in Fig S3(b). As a side note we would like to point that plotting the fluctuation spectrum as a function of **k*^2^*** as done by Straub and co-workers [41] forces the fit region (in this case the lower **k** modes) to imposes the region to artificially scale according to CWT. As seem in the fitting of the above discussed spectra, this need not be the case always.

It is important to note the different **k** regions fit to capillary wave theory in these two systems. Smaller **k** values scale according to CWT for the DPPC/TMP system whereas **k** values between 1 and 3 *nm*^−1^ scale according to CWT for DPPC/DUPC/CHOL system. The boundary length evolution data (see Fig. 3b) clearly suggests that even an 11 *µ*s of trajectory for the DPPC/DUPC/CHOL system is not sufficient to completely equilibrate the system. We also plot the spectra as a function of time for this system (see Fig. S5). The boundary length and line tension fitted to CWT regime is shown in the inset table. The changing spectra indicate that apart from the aforementioned reason for fitting higher **k** values due to inadequate sampling for complete equilibration in case of DPPC/DUPC/CHOL system, the scaling of fitting regions also depends on roughness of the boundary. Also, worth noting in the higher intensity of small wavelengths at the early stage of the domain formation when large frequency dominates the interface. To highlight this point, we artificially create a simulated roughness profiles of various frequency modes with different amplitudes (using a combination of sine waves) (see Fig. S6 - Fig. S9). For further clarity, we have included a discussion in section S3 of the SI about the nature of height fluctuations and the effect on the fluctuation spectra. For a rougher surface, more **k** modes show up with significant contribution to the fluctuation spectrum. Low **k** region of the fluctuation spectra obtained by Gaussian smoothing of the boundary scales with an *α* exponent value of 1.3, and fluctuation spectra of the molecularly rough boundary scales with *α* value of 0.04. From these two cases we observe that as the boundary gets smoother and looses the molecular roughness of its surface, the scaling according to CWT is better. The more intricate roughness of the boundary shows up as higher frequency modes in the fluctuation spectrum. Additional, to check of the deviation from fluctuation-dissipation in CWT is occurring due to the effect of dipole orientation on the interface Hamiltonian, we checked the orientation of the P-N vector with respect to the membrane normal (z-axis). As shown in Fig. S10 the orientation of the dipoles are very random for DPPC/DUPC/CHOL system. This could imply negligible correction to the line tension due to dipole orientations.

## 4 Conclusion

Using SVM classifier method to identify boundaries in phase separated CG and AA lipid systems, we have shown that this computationally less expensive method is efficient in locating simple and complex boundaries with a precision up to molecular length scales. Order parameters reported in literature including chemical identity of the lipids, tail order parameter *S_CC_* and degree of non-affinity in topological rearrangements of the lipids *χ*^2^ are used to distinguish *L_o_* and *L_d_* phases and fed into the SVM algorithm to determine boundary. We show that the difference in boundaries identified by each of these order parameters is reflected in the fluctuation spectra and line tension obtained by fitting the spectra.

These observations stress upon the sensitivity of line tension values to the microscopic fluctuations of the boundary. In experimental methods identification of the boundary is limited by resolution of the technique used. Lower resolutions of these methods in comparison to the inherent ability of molecular simulation trajectories to observe processes at molecular length scales causes blurring and artificial smoothing of the boundary fluctuations. Loss of molecular roughness of the boundary is reflected in its fluctuation spectrum. Depending on the roughness of the boundary, different regions of the fluctuation spectra scale as **k**^−2^ according to CWT. Fluctuations of a rougher boundary has significant contributions form the higher frequency modes in the fluctuation spectrum.

Apart from the dependence of scaling on the roughness of the boundary, we also find that the range of **k** values for which the fluctuation spectrum scales according to CWT depends on complete equilibration and statistically sufficient sampling of equilibrium state of the system. Based on the observations we hypothesize that absence of **k**^−2^ scaling for smaller **k** modes and the range of **k** values for which **k**^−2^ scaling is observed holds information about the non-thermal, active fluctuations in the system. In *in-vivo* set-ups constantly exposed to external perturbations, based on the criteria of range of **k** modes scaling according to CWT the equilibrium or active, out of equilibrium nature of the system can be studied.

## 5 Author Contributions

SSI and AS designed the research. SSI and AN implemented the algorithms, carried our programming and scripting and generated data. All authors analyzed the data and wrote the paper.

## 6 Acknowledgement

A.S. thanks Prof. Roland Netz and Prof. Prerna Sharma for their valuable comments and suggestions on the manuscript. The authors thank Peter Tieleman’s group at University of Calgary, Edward Lyman at University of Delaware and Alemayehu Gorfe at University of Texas Medical School for sharing trajectories of CG and AA systems used in this study. The authors thank the D.E. Shaw Research (Anton Supercomputer) for making available the bilayers molecular simulation trajectories from Edward Lyman’s group. A.S also thanks the faculty of Physics at IIT-Bombay (particularily Prof. Anirban Sain) for allowing Archit Negi to carry out his Bacherlor’s research work at IISc-Bangalore.

A.S. thanks the Indian Institute of Science and the Ministry of Human Resource Development of India for the start up grant and the Department of Science and Technology of India for the early career grant. This research was also supported by the Department of Biotechnology, Government of India in the form of IISc-DBT partnership programme. Support from FIST program sponsored by the Department of Science and Technology and UGC, India – Centre for Advanced Studies and Ministry of Human Resource Development, India is gratefully acknowledged by the authors.

## Supporting Information

### S1 Support Vector Machine algorithm

We have a set of training data with *n* points of form: (**x**_1_, *y*_1_), *…*(**x***_n_, y_n_*), where *y_i_* is either 1 or *−*1, giving the classification of the point **x_i_**, which is a *d* dimensional vector. The equation of any hyperplane is given by: **w^T^x** +*b* = 0 where **w** is a vector perpendicular to the hyperplane and **x** is any point on the hyperplane ^1^. If a hyperplane that successfully separates s called the separating hyperplane and a separating hyperplane that is farthest away from the points of either classes is the optimally separating hyperplane. The margin of a separating hyperplane is twice the distance between the hyperplane and the closest data point. The optimally separating hyperplane is the hyperplane which is equidistant from both of the classes and the vector points closest to the separating hyperplanes are called support vectors. The two parallel hyperplanes and the optimally separating hyper-plane are denoted as: **w^T^x** + *b* = *±*1 and **w^T^x** + *b* = 0. Since there are no points between the two hyperplanes, constrains for *y_i_ −*1 and +1 gives: *y_i_*(**w^T^x_i_** + *b*) ⩾ 1 ∀ 1 ⩽ *i* ⩽ *n*. The distance between any two parallel hyperplanes **w^T^x** + *b*_1_ = 0 and **w^T^x** + *b*_2_ = 0 is given by: 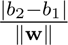. Therefore, the margin will be: 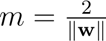. Maximizing *m*, or minimizing ||w|| with the constraint given above gives the optimally separating hyperplane. This essentially is an optimization problem and can be solved using the Lagrange multipliers method.

SVM is routinely used to classify linearly non-separable data. If we have data that is *d* dimensional, and is not linearly separable, we map it to a higher dimension where it is linearly separable and find the hyperplane in this dimension, and map it back to the initial dimension. The problem becomes computationally intractable with too many points or with highly increased dimensionality. Mapping every point to higher dimensions is not necessary, as the optimization problem requires only the pairwise dot products of all the points in the higher dimension. Kernel functions are used to do this and reduce the computation time considerably. Commonly used kernel functions are linear (*K*(**x**, **x**’) = **x**.**x**’), polynomial (*K*(**x**, **x**’) = (**x**.**x**’ + *c*)*^d^*), radial bias function (RBF, *K*(**x**, **x**’) = *exp*(*−γ*||**x** *−* **x**’||^2^)) and sigmoid (*K*(**x**, x’) = tanh (*κ***x**.**x**’ + *c*)) kernels.

### S2 Reconciling different formulations of LT calculation

Formulation 1 :

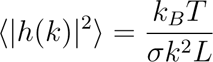

Formulation 2:

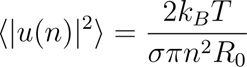

where 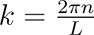, n is an integer, *σ* is line tension value, *R*_0_ is radius of the boundary, *L* is the boundary perimeter, *k_B_* is Boltzmann constant and *T* is temperature.

Hamiltonian of a fluctuating interface according to capillary wave theory is given by:

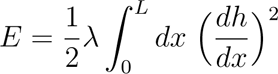

Here *h*(*x*) can be written as a Fourier series with periodicity [0,L]:

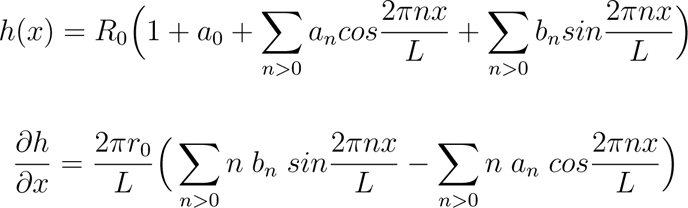

Here:

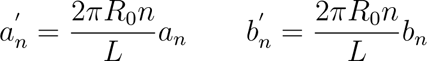

Using Parseval’s identity (given) in energy equation:

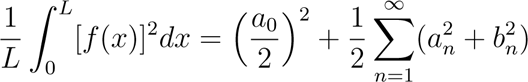

where *a*_0_ = 0. Hence:

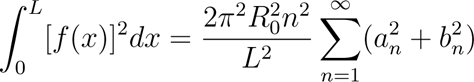

Substituting in energy equation:

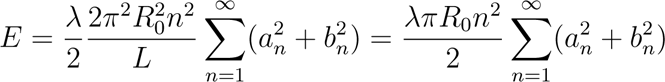

Applying equipartition theorem which states that each mode contributes 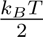 energy:

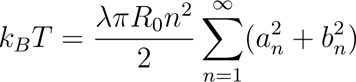

Hence:

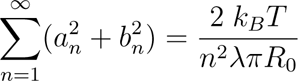

Thus, we arrive at the expression for formalism 2^2^ starting from Hamiltonian used in formalism 1^3^.

Another way of showing equivalence of the two formulations is shown below: Writing *h*(*x*) as a Fourier series in *x* similar to ref^?^ :

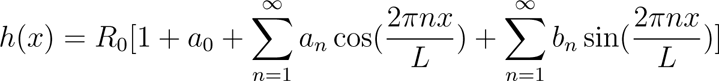

Here, *a_n_* and *b_n_* are dimensionless, and *R*_0_ has dimension of length, 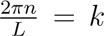 and the coefficients of the *n^th^* Fourier series terms are *R*_0_*a_n_* for cos and *R*_0_*b_n_* for sin terms. According to Sarah Keller and co-workers ^3^:

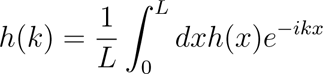

However, at integer indices, *h*(*k*) will be related to the Fourier series coefficients as:

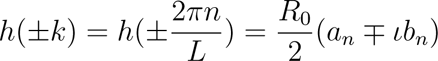

The *R*_0_ is present in the numerator as the coefficients in the Fourier series are *R*_0_*a_n_* and *R*_0_*b_n_* instead of *a_n_* and *b_n_*. Therefore:

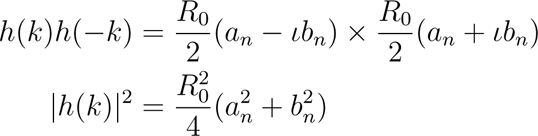

The two formulations we have are:

Formulation 1 :

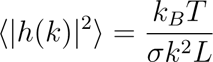

Formulation 2:

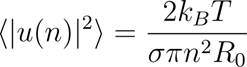

where 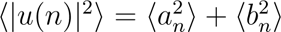 (without the *R*_0_ factor, as that is how it was in the ref **^?^**), 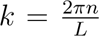, n is an integer, *σ* is line tension value, *R*_0_ is radius of the boundary, *L* is the boundary perimeter, *k_B_* is Boltzmann constant and *T* is temperature.

Proceeding from formulation 1:

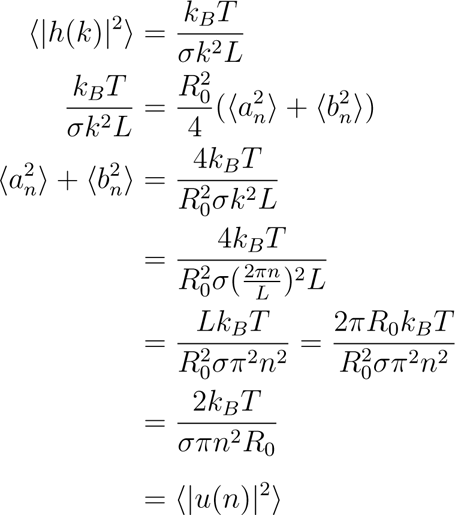

Therefore, formulation 2 follows from formulation 1 and vice versa.

**Figure S1:**
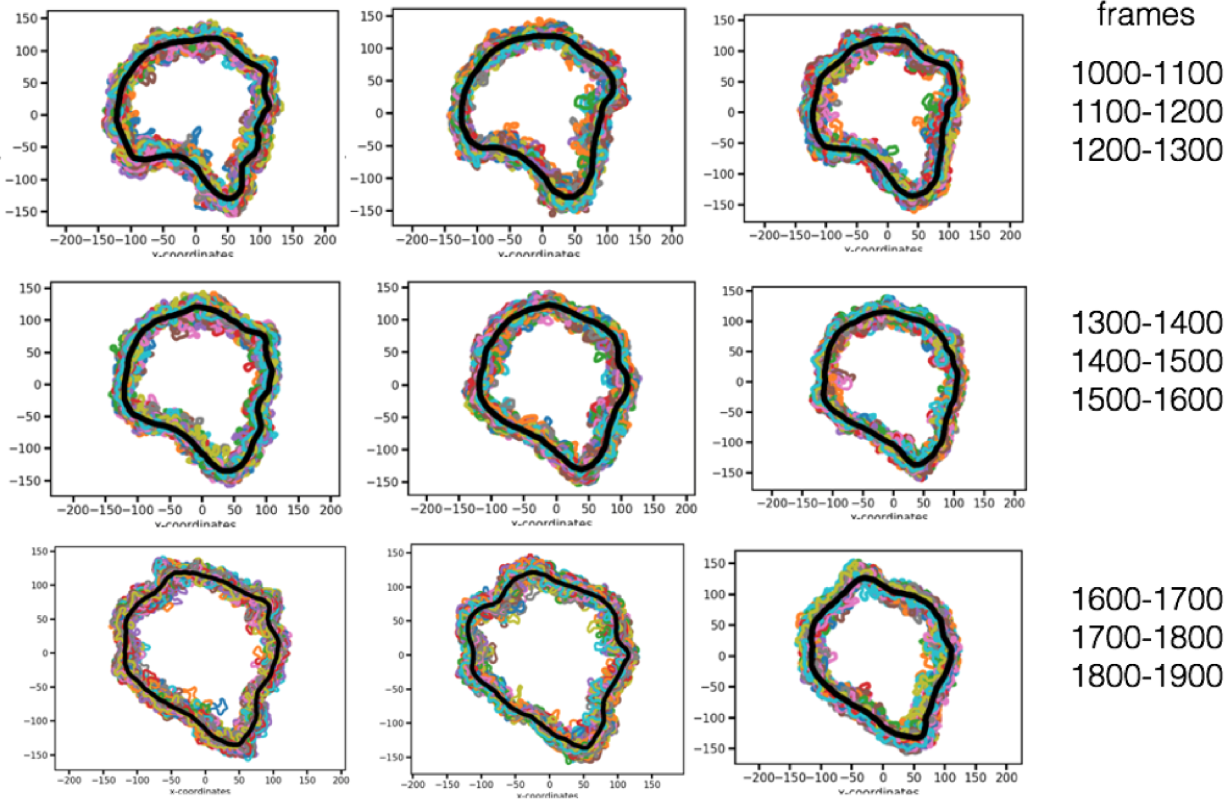
Boundaries identified based on their chemical identity and their average shown in black for three independent fragments of trajectory of the DPPC/DUPC/CHOL system. Every independent trajectory contains 100 frames.

**Figure S2:**
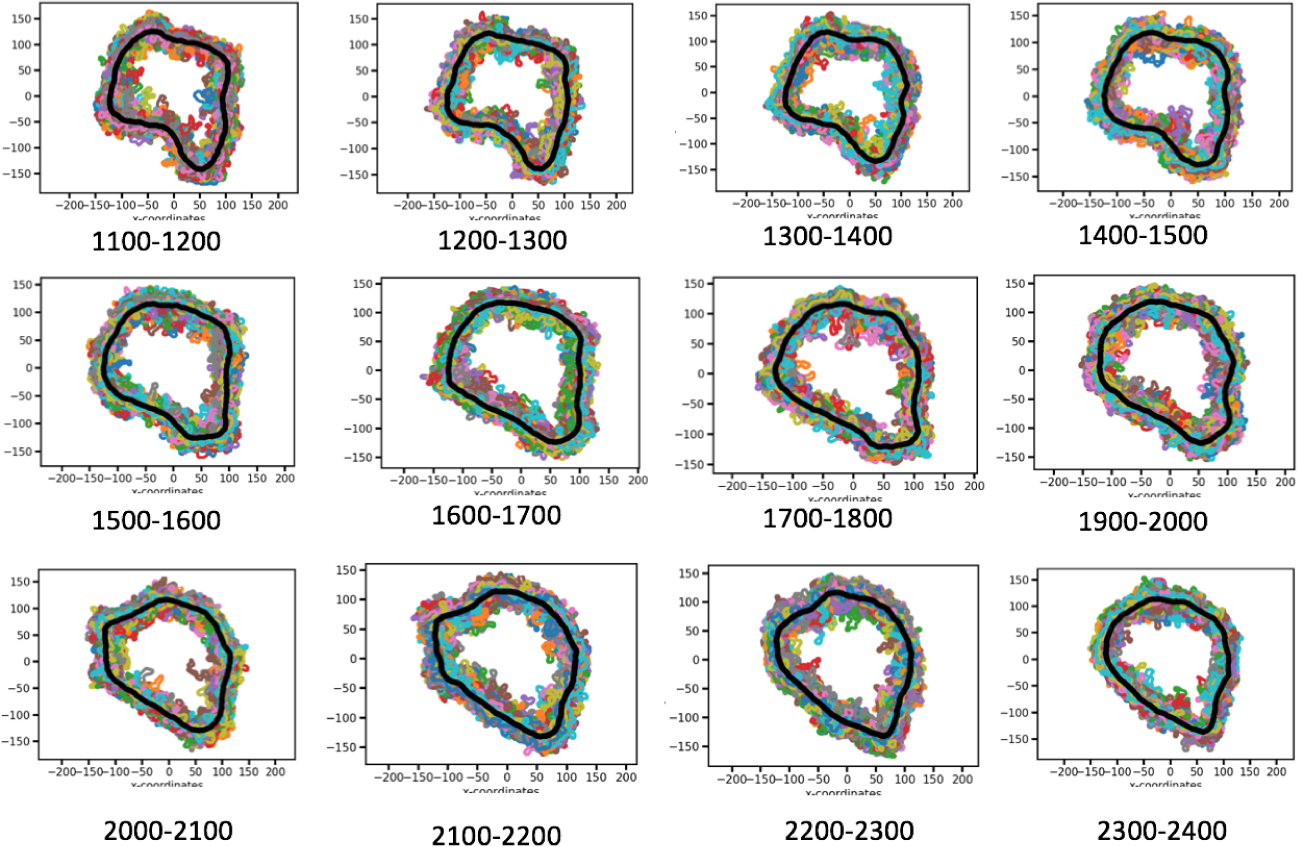
Boundaries identified based on on a cut-off of 0.4 on the *S_CC_* values and their average shown in black for three independent fragments of trajectory of the DPPC/DUPC/CHOL system. Every independent trajectory contains 100 frames.

**Figure S3:**
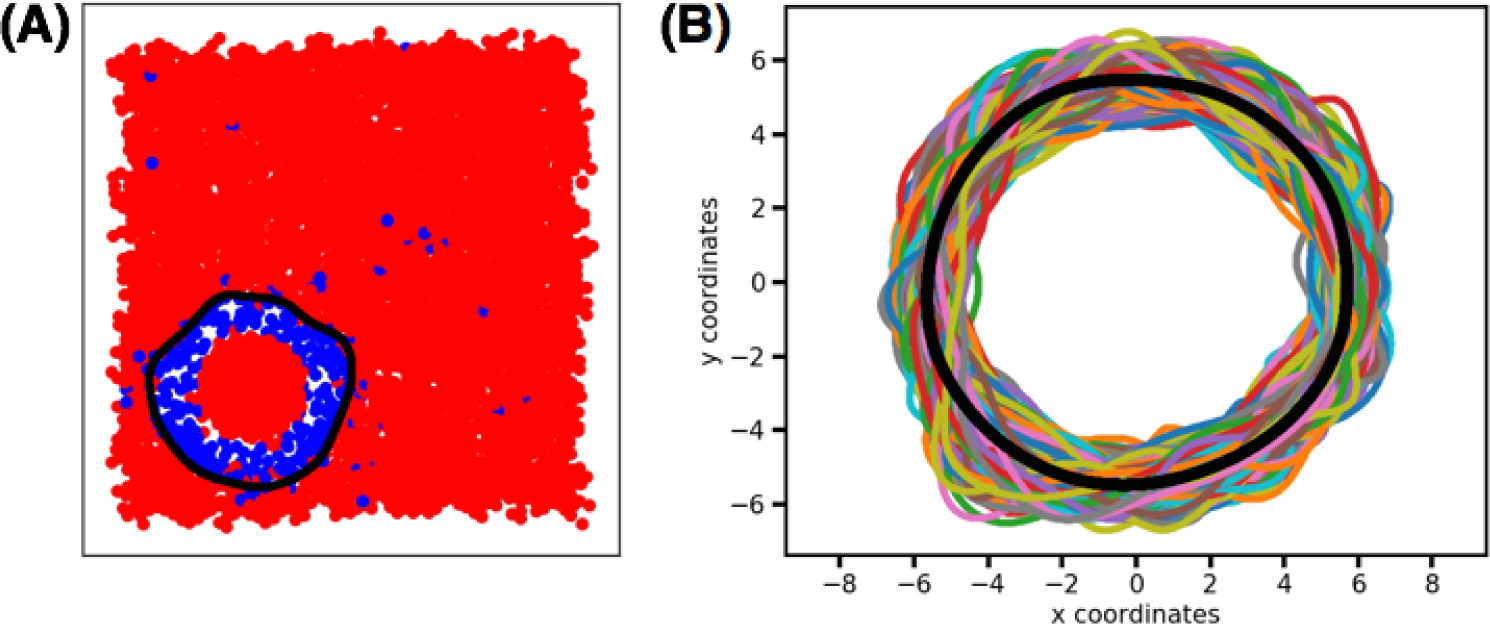
Boundaries identified based a cut-off of 2000 on the *χ*^2^ values and their average shown in black for three independent fragments of trajectory of the DPPC/DUPC/CHOL system. Each boundary shown is averaged every 5 frames. Hence the 100 boundaries shown in the figure corresponds to 500 frames.

**Figure S4:**
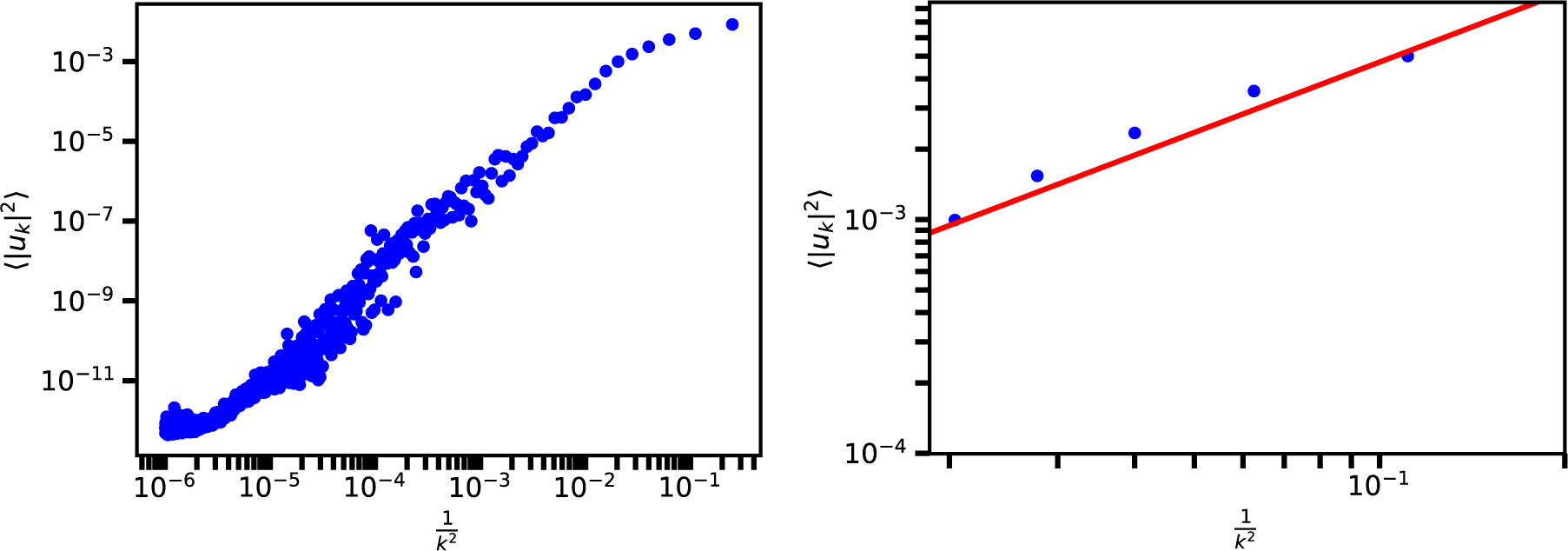
Spectra of boundary fluctuation for DPPC/TMP system using formulation 2 (by Tobias Baugmart et al). Figure on the left shows the full fluctuation spectrum and on the right the region fit to capillary wave theory to obtain line tension is shown. The fitting gives a line tension value of 10.01 pN.

**Figure S5:**
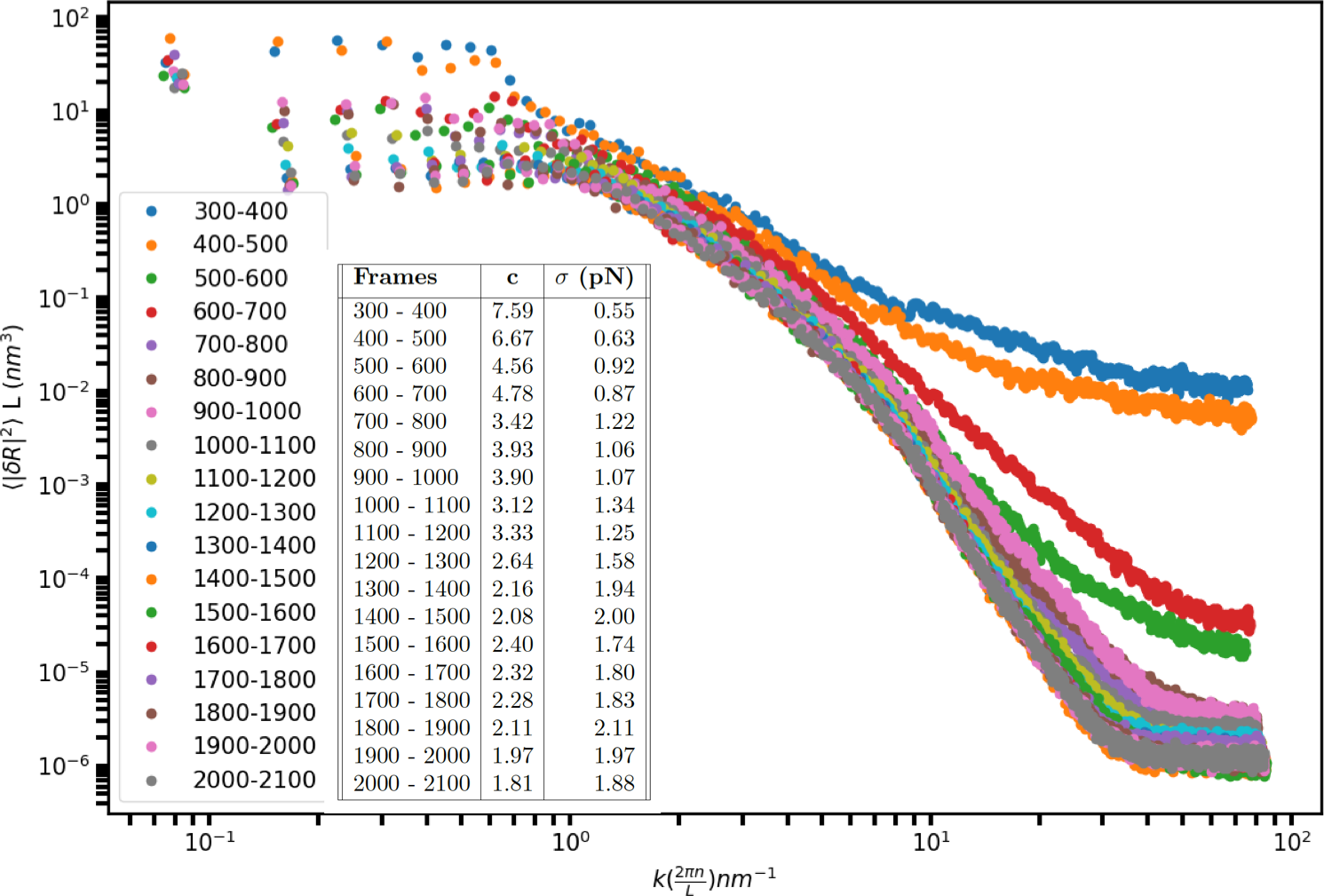
Spectra of boundary fluctuation for DPPC/DUPC/CHOL system for different frame ranges. As the system evolves, slope of the region between k=1 to 6 decreases.

### S3 Simulated hight fluctuations and their spectra - understanding the fluctuation spectra

In this section we discuss fluctuation spectra of simulated height fluctuations generated by a combination of sine waves. Figure S6 shows the fluctuation spectrum of height fluctuations consisting of low amplitude, small frequency vibrations. Fluctuation spectrum for systems with increasing higher frequencies and with higher amplitudes are shown in Figure S8 and Figure S9 respectively. Higher frequency and very large amplitude fluctuations are less commonly observed in biological systems. These fluctuation violate equipartition theory. For fluctuation spectrum shown in Figure S6, Figure S7 shows how the height fluctuations are reconstructed from the different mode contributions given by the spectrum analysis.

The differences in the profiles of the fluctuation spectrum for each of these cases shows that the region of fluctuation spectrum that carries useful information about the boundary (height) fluctuations differs according to amplitudes and frequencies of these fluctuations.

**Figure S6:**
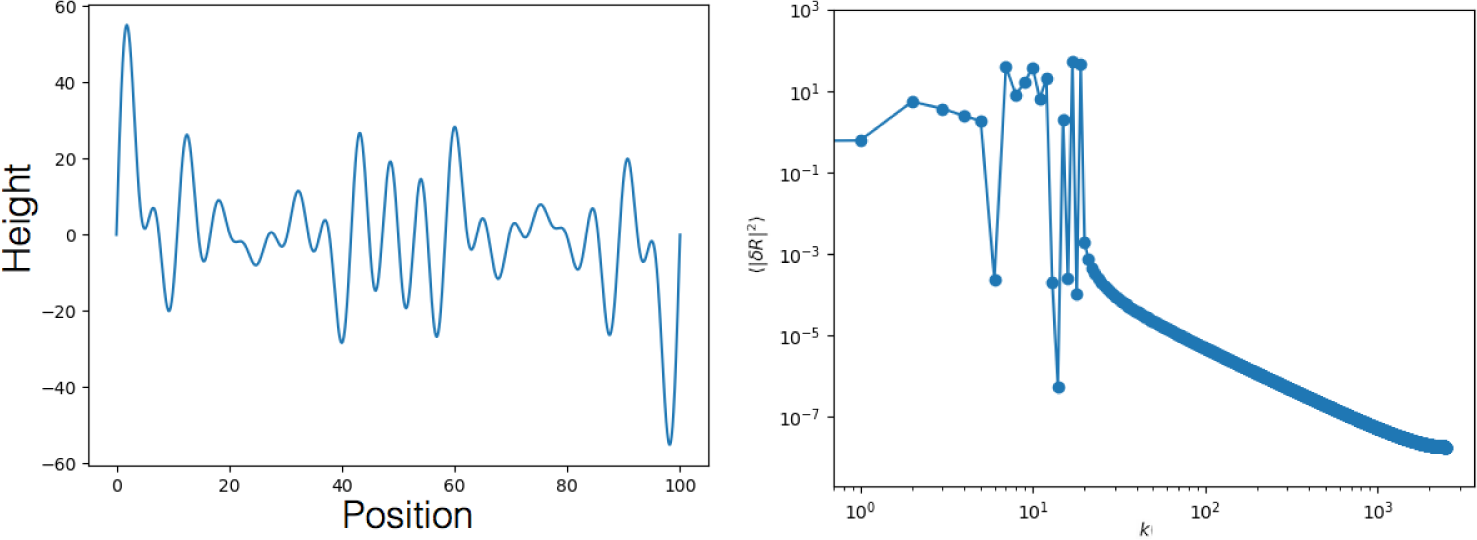
Simulated height fluctuations using 14 sine waves of different amplitude and frequency (left) and its corresponding fluctuation spectrum (right).

**Figure S7:**
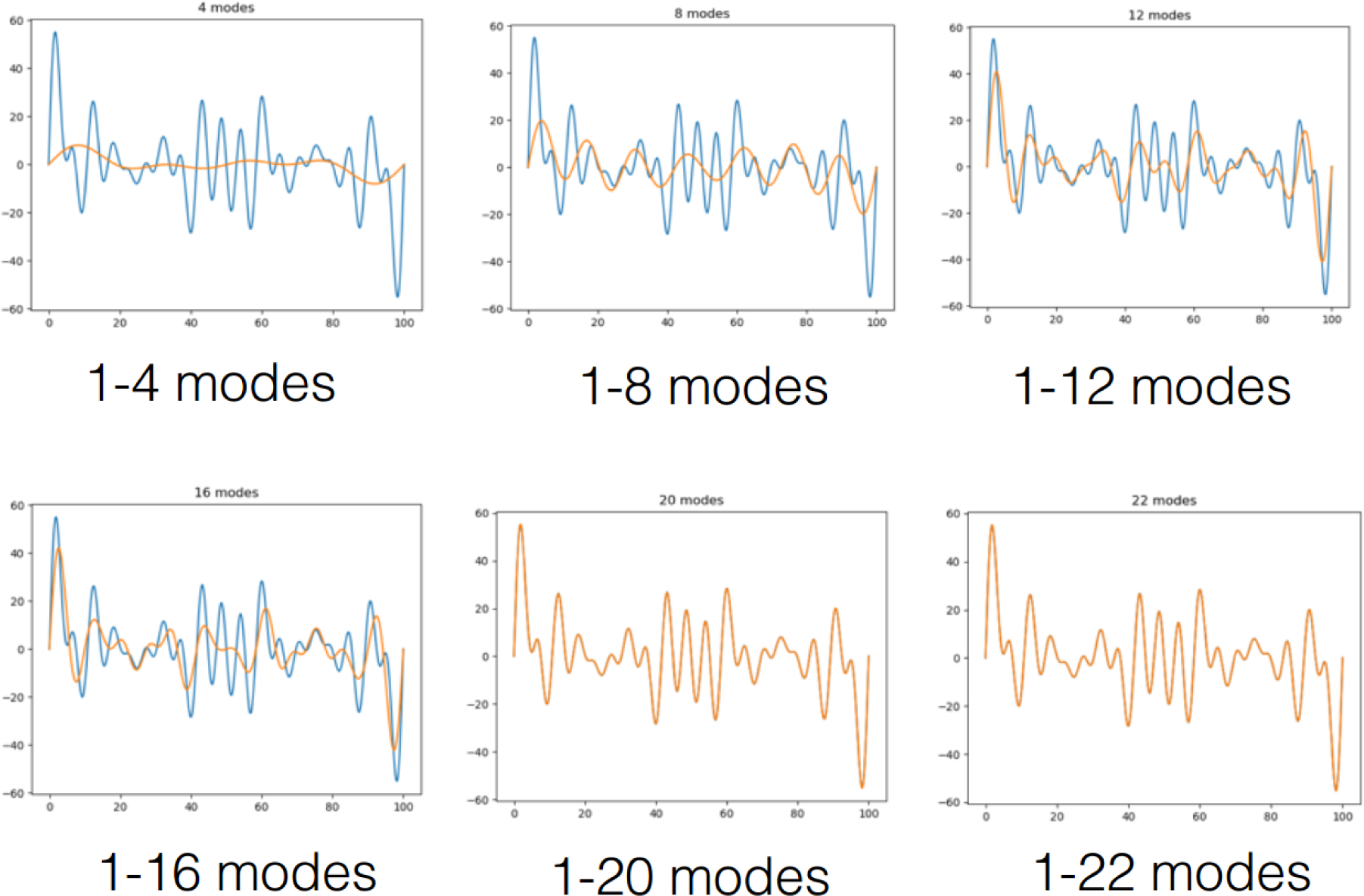
Simulated height fluctuations using 22 sine waves of different amplitude and higher frequencies shown in blue. The wave reconstructed with different mode regions is shown as a overlay on the blue curve. First 20 modes are enough to fully capture the fluctuations (shown by a perfect overlap of blue and orange curves).

**Figure S8:**
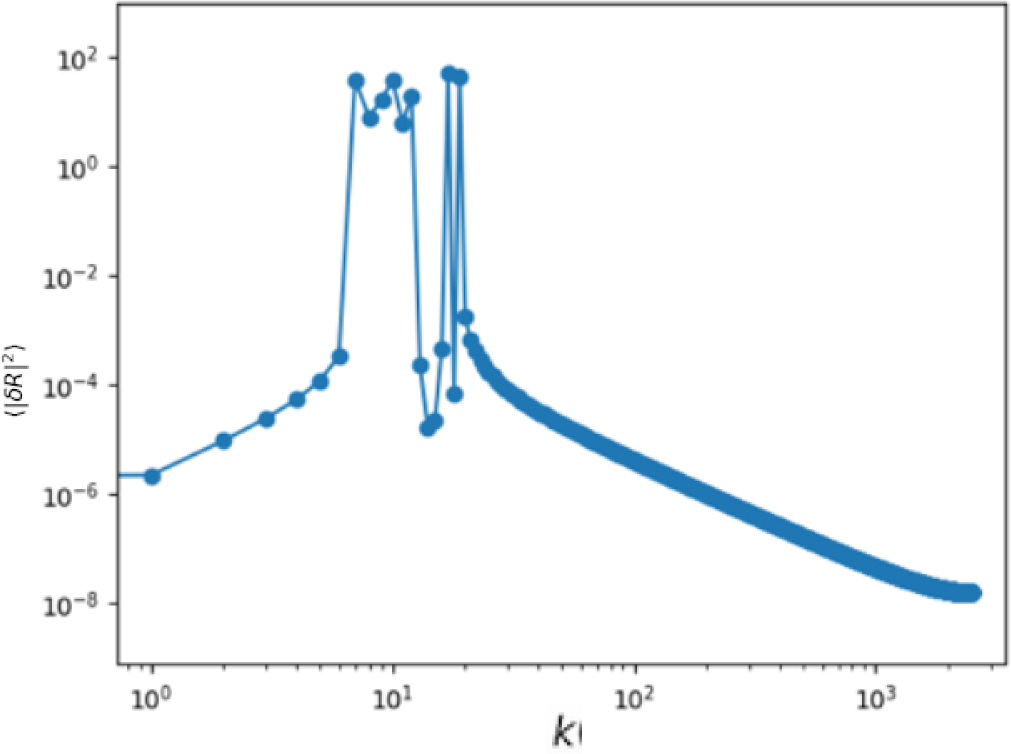
Spectrum corresponding to heigh fluctuations shown in Figure S7. Contribution of low k region in negligible due to the construction of the simulated hight fluctuations with higher frequencies.

**Figure S9:**
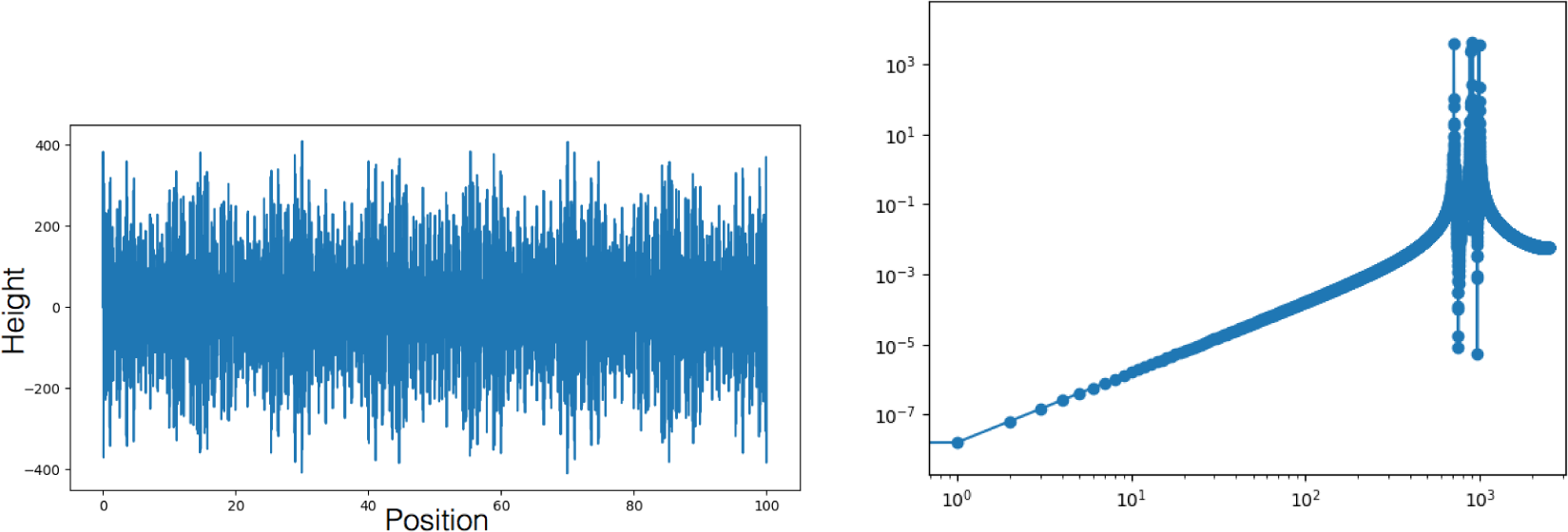
Simulated height fluctuations using high amplitude amplitude, high frequency sine waves (left) and its corresponding fluctuation spectrum (right). The fluctuation spectrum looks like the mirror image of spectrum shown in Figure S6.

**Figure S10:**
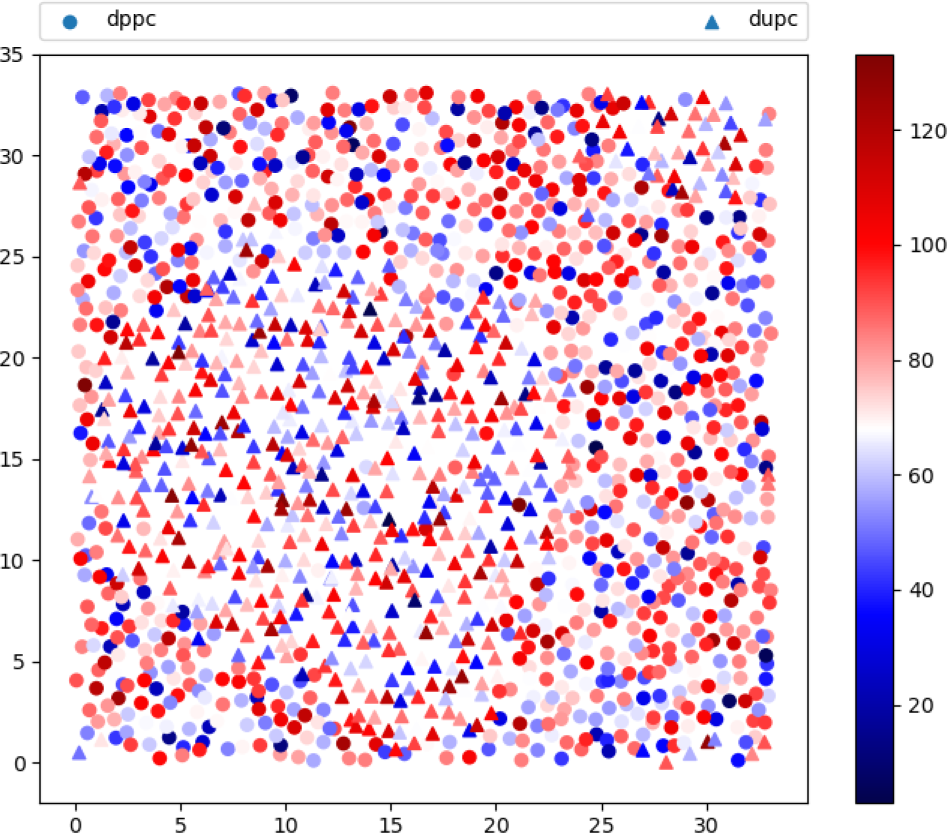
Random orientation of P-N vector with respect to the membrane normal for DPPC/DUPC/CHOL system.

### S4 Movies

movS1a.mov - Movie showing one of the boundaries in DAPC/DPPC/CHOL system. *L_o_*(blue) and *L_d_* (red) are marked based on chemical identity.

movS1b.mov - Movie showing one of the boundaries in DAPC/DPPC/CHOL system with tLAT peptide localizing at boundary interface. *L_o_* (blue) and *L_d_* (red) are marked based on chemical identity. Peptides are shown in purple.

movS2a.mov - Movie showing phase separation in DUPC/DPPC/CHOL system monitored using chemical identity. Ld phase is shown in blue and Lo phase in red.

movS2b.mov - Boundary identified using chemical identity of lipids shown for representative 100 snapshots. DUPC - red, DPPC - blue, boundary black.

movS3a.mov - Movie showing phase separation in DUPC/DPPC/CHOL system monitored using *S_cc_* values.

movS3b.mov - Movie showing phase separation in DUPC/DPPC/CHOL system with *L_d_* in red and *L_o_* in blue. *L_o_* and *L_d_* are decided using a cut-off of 0.5 on the *S_cc_* values.

movS3c.mov - Movie showing boundary marked using a cut-off on *S_cc_* values for last 100 frames. *L_d_* shown in red and *L_o_* shown in blue. *L_o_* and *L_d_* are decided using a cut-off of 0.5 on the *S_cc_* values.

movS4a.mov - Movie showing phase separation in DUPC/DPPC/CHOL system monitored using *χ*^2^ values.

movS4b.mov - Movie showing phase separation in DUPC/DPPC/CHOL system monitored coloured as Lo (red) and Ld (blue) based on cut-off of 2000 of *χ*^2^ values.

movS4c.mov - Movie showing boundary marked using a cut-off on *χ*^2^ values averaged every 5 frames. *L_d_* shown in red and *L_o_* shown in blue. *L_o_* and *L_d_* are decided using a cut-off of 2000 on the *χ*^2^ values.

MovS5.mov - Movie showing boundary in the DPPC/TMP system. Boundary marked in black is identified using a cut-off on *χ*^2^ values.

MovS6a.mov - Movie showing DPPC/DUPC/CHOL grid points coloured based on thickness values - monitored for the last 100 frames.

MovS6b.mov - Movie showing DPPC/DUPC/CHOL grid points coloured based on cut-off of 4.3 nm applied on thickness values - monitored for the last 100 frames. Grid points above 4.3 nm cut-off are shown in blue and below 4.3 nm are shown in red.

MovS7a.mov - Movie showing DPPC/DUPC/CHOL grid points coloured based on *ϕ*_6_ values - monitored for the last 100 frames.

MovS7b.mov - Movie showing DPPC/DUPC/CHOL sites coloured based on cut-off of 0.2 applied on *ϕ*_6_ values - monitored for the last 100 frames. Sites above 0.2 cut-off are shown in red and below 0.2 nm are shown in blue. Boundary obtained using SVM is shown in black.

## References

[1] E. J. Shimshick and H. M. McConnell, “Lateral phase separation in phospholipid membranes,” Biochemistry, vol. 12, no. 12, pp. 2351–2360, 1973.

[2] J. Seelig and W. Niederberger, “Two pictures of a lipid bilayer. comparison between deuterium label and spin-label experiments,” Biochemistry, vol. 13, no. 8, pp. 1585–1588, 1974.

[3] V. A. Petit and M. Edidin, “Lateral phase separation of lipids in plasma membranes: effect of temperature on the mobility of membrane antigens,” Science, vol. 184, no. 4142, pp. 1183–1185, 1974.

[4] A. Seelig and J. Seelig, “Dynamic structure of fatty acyl chains in a phospholipid bilayer measured by deuterium magnetic resonance,” Biochemistry, vol. 13, no. 23, pp. 4839–4845, 1974.

[5] A. Helenius and K. Simons, “Solubilization of membranes by detergents,” Biochimica et Biophysica Acta (BBA) - Reviews on Biomembranes, vol. 415, no. 1, pp. 29–79, 1975.

[6] A. Lee, N. Birdsall, J. Metcalfe, G. Warren, and G. Roberts, “A determination of the mobility gradient in lipid bilayers by 13 c nuclear magnetic resonance,” Proceedings of the Royal Society of London - Biological Sciences, vol. 193, no. 1112, pp. 253–274, 1976.

[7] J. H. Ipsen, G. Karlström, O. Mourtisen, H. Wennerström, and M. Zuckermann, “Phase equilibria in the phosphatidylcholine-cholesterol system,” Biochimica et Biophysica Acta (BBA) - Biomembranes, vol. 905, no. 1, pp. 162–172, 1987.

[8] H. M. McConnell and V. T. Moy, “Shapes of finite two-dimensional lipid domains,” The Journal of Physical Chemistry, vol. 92, no. 15, pp. 4520–4525, 1988.

[9] P. F. F. Almeida, W. L. C. Vaz, and T. E. Thompson, “Lateral diffusion in the liquid phases of dimyristoylphosphatidylcholine/cholesterol lipid bilayers: a free volume analysis,” Biochemistry, vol. 31, no. 29, pp. 6739–6747, 1992.

[10] D. J. Benvegnu and H. M. McConnell, “Line tension between liquid domains in lipid monolayers,” The Journal of Physical Chemistry, vol. 96, no. 16, pp. 6820–6824, 1992.

[11] C. Dietrich, L. Bagatolli, Z. Volovyk, N. Thompson, M. Levi, K. Jacobson, and E. Gratton, “Lipid rafts reconstituted in model membranes,” Biophysical journal, vol. 80, no. 3, pp. 1417–1428, 2001.

[12] S. L. Veatch and S. L. Keller, “Organization in lipid membranes containing cholesterol,” Physical review letters, vol. 89, no. 26, p. 268101, 2002.

[13] S. L. Veatch and S. L. Keller, “Separation of liquid phases in giant vesicles of ternary mixtures of phospholipids and cholesterol,” Biophysical journal, vol. 85, no. 5, pp. 3074–3083, 2003.

[14] D. Marsh, “Cholesterol-induced fluid membrane domains: a compendium of lipidraft ternary phase diagrams,” Biochimica et Biophysica Acta (BBA)-Biomembranes, vol. 1788, no. 10, pp. 2114–2123, 2009.

[15] F. A. Heberle and G. W. Feigenson, “Phase separation in lipid membranes,” Cold Spring Harbor perspectives in biology, vol. 3, no. 4, p. a004630, 2011.

[16] D. G. Ackerman and G. W. Feigenson, “Multiscale modeling of four-component lipid mixtures: domain composition, size, alignment, and properties of the phase interface,” The Journal of Physical Chemistry B, vol. 119, no. 11, pp. 4240–4250, 2015.

[17] S. L. Goh, J. J. Amazon, and G. W. Feigenson, “Toward a better raft model: modulated phases in the four-component bilayer, dspc/dopc/popc/chol,” Biophysical journal, vol. 104, no. 4, pp. 853–862, 2013.

[18] T. M. Konyakhina, J. Wu, J. D. Mastroianni, F. A. Heberle, and G. W. Feigenson, “Phase diagram of a 4-component lipid mixture: Dspc/dopc/popc/chol,” Biochimica et Biophysica Acta (BBA)-Biomembranes, vol. 1828, no. 9, pp. 2204–2214, 2013.

[19] H.-M. Wu, Y.-H. Lin, T.-C. Yen, and C.-L. Hsieh, “Nanoscopic substructures of raft-mimetic liquid-ordered membrane domains revealed by high-speed single-particle tracking,” Scientific reports, vol. 6, p. 20542, 2016.

[20] H. Mizuno, M. Abe, P. Dedecker, A. Makino, S. Rocha, Y. Ohno-Iwashita, J. Hofkens, T. Kobayashi, and A. Miyawaki, “Fluorescent probes for superresolution imaging of lipid domains on the plasma membrane,” Chemical Science, vol. 2, no. 8, pp. 1548–1553, 2011.

[21] M. B. Stone, S. A. Shelby, and S. L. Veatch, “Super-resolution microscopy: shedding light on the cellular plasma membrane,” Chemical reviews, vol. 117, no. 11, pp. 7457–7477, 2017.

[22] D. M. Owen, D. J. Williamson, A. Magenau, and K. Gaus, “Sub-resolution lipid domains exist in the plasma membrane and regulate protein diffusion and distribution,” Nature communications, vol. 3, p. 1256, 2012.

[23] S. Rayermann, G. Rayermann, A. Merz, and S. Keller, “Hallmarks of reversible phase separation in living, unperturbed cell membranes,” Biophysical Journal, vol. 112, no. 3, Supplement 1, p. 522a, 2017.

[24] R. F. De Almeida, A. Fedorov, and M. Prieto, “Sphingomyelin/phosphatidylcholine/cholesterol phase diagram: boundaries and composition of lipid rafts,” Biophysical journal, vol. 85, no. 4, pp. 2406–2416, 2003.

[25] A. J. García-Sáez, S. Chiantia, and P. Schwille, “Effect of line tension on the lateral organization of lipid membranes,” Journal of Biological Chemistry, vol. 282, no. 46, pp. 33537–33544, 2007.

[26] R. D. Usery, T. A. Enoki, S. P. Wickramasinghe, M. D. Weiner, W.-C. Tsai, M. B. Kim, S. Wang, T. L. Torng, D. G. Ackerman, F. A. Heberle, et al., “Line tension controls liquid-disordered+ liquid-ordered domain size transition in lipid bilayers,” Biophysical journal, vol. 112, no. 7, pp. 1431–1443, 2017.

[27] J. W. Gibbs, “Art. lii.–on the equilibrium of heterogeneous substances,” American Journal of Science and Arts (1820-1879), vol. 16, no. 96, p. 441, 1878.

[28] J. Gibbs, “The collected papers of j. willard gibbs,” Yale University Press, London, vol. 1, pp. 55–353, 1957.

[29] F. A. Heberle, R. S. Petruzielo, J. Pan, P. Drazba, N. Kučerka, R. F. Standaert, G. W. Feigenson, and J. Katsaras, “Bilayer thickness mismatch controls domain size in model membranes,” Journal of the American Chemical Society, vol. 135, no. 18, pp. 6853–6859, 2013.

[30] C. M. Rosetti, G. G. Montich, and C. Pastorino, “Molecular insight into the line tension of bilayer membranes containing hybrid polyunsaturated lipids,” The Journal of Physical Chemistry B, vol. 121, no. 7, pp. 1587–1600, 2017.

[31] S. S. Iyer and A. Srivastava, “Intrinsic disorder and degeneracy in molecular scale organization of biological membrane,” bioRxiv, 2019.

[32] S. Safran, Statistical thermodynamics of surfaces, interfaces, and membranes. CRC Press, 2018.

[33] J. A. Killian, “Hydrophobic mismatch between proteins and lipids in membranes,” Biochimica et Biophysica Acta (BBA)-Reviews on Biomembranes, vol. 1376, no. 3, pp. 401–416, 1998.

[34] O. Mouritsen and M. Bloom, “Mattress model of lipid-protein interactions in membranes,” Biophysical journal, vol. 46, no. 2, pp. 141–153, 1984.

[35] S. Katira, K. K. Mandadapu, S. Vaikuntanathan, B. Smit, and D. Chandler, “Pre-transition effects mediate forces of assembly between transmembrane proteins,” Elife, vol. 5, p. e13150, 2016.

[36] C. D. Blanchette, W.-C. Lin, C. A. Orme, T. V. Ratto, and M. L. Longo, “Using nucleation rates to determine the interfacial line tension of symmetric and asymmetric lipid bilayer domains,” Langmuir, vol. 23, no. 11, pp. 5875–5877, 2007.

[37] A. R. Honerkamp-Smith, P. Cicuta, M. D. Collins, S. L. Veatch, M. den Nijs, M. Schick, and S. L. Keller, “Line tensions, correlation lengths, and critical exponents in lipid membranes near critical points,” Biophysical journal, vol. 95, no. 1, pp. 236–246, 2008.

[38] C. Esposito, A. Tian, S. Melamed, C. Johnson, S.-Y. Tee, and T. Baumgart, “Flicker spectroscopy of thermal lipid bilayer domain boundary fluctuations,” Biophysical journal, vol. 93, no. 9, pp. 3169–3181, 2007.

[39] A. Tian, C. Johnson, W. Wang, and T. Baumgart, “Line tension at fluid membrane domain boundaries measured by micropipette aspiration,” Physical review letters, vol. 98, no. 20, p. 208102, 2007.

[40] J. B. Hutchison, R. M. Weis, and A. D. Dinsmore, “Change of line tension in phase-separated vesicles upon protein binding,” Langmuir, vol. 28, no. 11, pp. 5176–5181, 2012.

[41] A. Bandara, A. Panahi, G. A. Pantelopulos, T. Nagai, and J. E. Straub, “Exploring the impact of proteins on the line tension of a phase-separating ternary lipid mixture,” The Journal of chemical physics, vol. 150, no. 20, p. 204702, 2019.

[42] A. J. Sodt, R. W. Pastor, and E. Lyman, “Hexagonal substructure and hydrogen bonding in liquid-ordered phases containing palmitoyl sphingomyelin,” Biophysical journal, vol. 109, no. 5, pp. 948–955, 2015.

[43] A. J. Sodt, M. L. Sandar, K. Gawrisch, R. W. Pastor, and E. Lyman, “The molecular structure of the liquid-ordered phase of lipid bilayers,” Journal of the American Chemical Society, vol. 136, no. 2, pp. 725–732, 2014.

[44] S. J. Marrink, H. J. Risselada, S. Yefimov, D. P. Tieleman, and A. H. De Vries, “The martini force field: coarse grained model for biomolecular simulations,” The journal of physical chemistry B, vol. 111, no. 27, pp. 7812–7824, 2007.

[45] S. Baoukina, E. Mendez-Villuendas, W. D. Bennett, and D. P. Tieleman, “Computer simulations of the phase separation in model membranes,” Faraday discussions, vol. 161, pp. 63–75, 2013.

[46] X. Lin, A. A. Gorfe, and I. Levental, “Protein partitioning into ordered membrane domains: insights from simulations,” Biophysical journal, vol. 114, no. 8, pp. 1936–1944, 2018.

[47] T. J. Piggot, J. R. Allison, R. B. Sessions, and J. W. Essex, “On the calculation of acyl chain order parameters from lipid simulations,” Journal of chemical theory and computation, vol. 13, no. 11, pp. 5683–5696, 2017.

[48] N. D. Mermin, “Crystalline order in two dimensions,” Physical Review, vol. 176, no. 1, p. 250, 1968.

[49] M. L. Falk and J. S. Langer, “Dynamics of viscoplastic deformation in amorphous solids,” Physical Review E, vol. 57, no. 6, p. 7192, 1998.

[50] S. S. Iyer, M. Tripathy, and A. Srivastava, “Fluid phase coexistence in biological membrane: Insights from local nonaffine deformation of lipids,” Biophysical journal, vol. 115, no. 1, pp. 117–128, 2018.

[51] M. Tripathy, S. S. Iyer, and A. Srivastava, “Molecular origin of spatiotemporal heterogeneity in biomembranes with coexisting liquid phases: Insights from topological rearrangements and lipid packing defects,” in Advances in Biomembranes and Lipid Self-Assembly, vol. 28, pp. 87–114, Elsevier, 2018.

[52] F. Buff, R. Lovett, and F. Stillinger Jr, “Interfacial density profile for fluids in the critical region,” Physical Review Letters, vol. 15, no. 15, p. 621, 1965.

[53] M. Jorge and M. N. D. Cordeiro, “Intrinsic structure and dynamics of the water/nitrobenzene interface,” The Journal of Physical Chemistry C, vol. 111, no. 47, pp. 17612–17626, 2007.

[54] A. P. Willard and D. Chandler, “Instantaneous liquid interfaces,” The Journal of Physical Chemistry B, vol. 114, no. 5, pp. 1954–1958, 2010.

[55] B. E. Boser, I. M. Guyon, and V. N. Vapnik, “A training algorithm for optimal margin classifiers,” in Proceedings of the fifth annual workshop on Computational learning theory, pp. 144–152, ACM, 1992.

[56] V. N. Vapnik, “An overview of statistical learning theory,” IEEE transactions on neural networks, vol. 10, no. 5, pp. 988–999, 1999.

[57] S. Hua and Z. Sun, “A novel method of protein secondary structure prediction with high segment overlap measure: support vector machine approach,” Journal of molecular biology, vol. 308, no. 2, pp. 397–407, 2001.

[58] C. Cai, L. Han, Z. L. Ji, X. Chen, and Y. Z. Chen, “Svm-prot: web-based support vector machine software for functional classification of a protein from its primary sequence,” Nucleic acids research, vol. 31, no. 13, pp. 3692–3697, 2003.

[59] C. H. Ding and I. Dubchak, “Multi-class protein fold recognition using support vector machines and neural networks,” Bioinformatics, vol. 17, no. 4, pp. 349–358, 2001.

[60] S. J. Irausquin and L. Wang, “A machine learning approach for prediction of lipid-interacting residues in amino acid sequences,” in 2007 IEEE 7th International Symposium on BioInformatics and BioEngineering, pp. 315–319, IEEE, 2007.

[61] E. Jones, T. Oliphant, P. Peterson, et al., “SciPy: Open source scientific tools for Python,” 2001–.

[62] F. Pedregosa, G. Varoquaux, A. Gramfort, V. Michel, B. Thirion, O. Grisel, M. Blondel, P. Prettenhofer, R. Weiss, V. Dubourg, J. Vanderplas, A. Passos, D. Cournapeau, M. Brucher, M. Perrot, and E. Duchesnay, “Scikit-learn: Machine learning in Python,” Journal of Machine Learning Research, vol. 12, pp. 2825–2830, 2011.

[63] H. T. Davis, “Capillary waves and the mean field theory of interfaces,” The Journal of Chemical Physics, vol. 67, no. 8, pp. 3636–3641, 1977.

[64] J. S. Rowlinson and B. Widom, Molecular theory of capillarity. Courier Corporation, 2013.

[65] D. Bedeaux and J. D. Weeks, “Correlation functions in the capillary wave model of the liquid–vapor interface,” The Journal of chemical physics, vol. 82, no. 2, pp. 972–979, 1985.

[66] B. A. Camley and F. L. H. Brown, “Dynamic simulations of multicomponent lipid membranes over long length and time scales,” Phys. Rev. Lett., vol. 105, p. 148102, Sep 2010.

[67] B. A. Camley, C. Esposito, T. Baumgart, and F. L. Brown, “Lipid bilayer domain fluctuations as a probe of membrane viscosity,” Biophysical Journal, vol. 99, no. 6, pp. L44–L46, 2010.

[68] S. Baoukina, E. Mendez-Villuendas, and D. P. Tieleman, “Molecular view of phase coexistence in lipid monolayers,” Journal of the American Chemical Society, vol. 134, no. 42, pp. 17543–17553, 2012.

[69] P. Tarazona and E. Chacón, “Monte carlo intrinsic surfaces and density profiles for liquid surfaces,” Physical Review B, vol. 70, no. 23, p. 235407, 2004.

[70] W. J. Allen, J. A. Lemkul, and D. R. Bevan, “Gridmat-md: a grid-based membrane analysis tool for use with molecular dynamics,” Journal of computational chemistry, vol. 30, no. 12, pp. 1952–1958, 2009.

[71] H. Strey, M. Peterson, and E. Sackmann, “Measurement of erythrocyte membrane elasticity by flicker eigenmode decomposition,” Biophysical Journal, vol. 69, no. 2, pp. 478–488, 1995.

[72] H.-G. Döbereiner, G. Gompper, C. K. Haluska, D. M. Kroll, P. G. Petrov, and K. A. Riske, “Advanced flicker spectroscopy of fluid membranes,” Physical review letters, vol. 91, no. 4, p. 048301, 2003.

[73] J. Pécréaux, H.-G. Döbereiner, J. Prost, J.-F. Joanny, and P. Bassereau, “Refined contour analysis of giant unilamellar vesicles,” The European Physical Journal E, vol. 13, no. 3, pp. 277–290, 2004.

[74] C. Monzel and K. Sengupta, “Measuring shape fluctuations in biological membranes,” Journal of Physics D: Applied Physics, vol. 49, no. 24, p. 243002, 2016.

## References

[1] A. Kowalczyk, “Support vector machines succinctly,” Syncfusion Inc, 2017.

[2] C. Esposito, A. Tian, S. Melamed, C. Johnson, S.-Y. Tee, and T. Baumgart, “Flicker spectroscopy of thermal lipid bilayer domain boundary fluctuations,” Biophysical journal, vol. 93, no. 9, pp. 3169–3181, 2007.

[3] A. R. Honerkamp-Smith, P. Cicuta, M. D. Collins, S. L. Veatch, M. den Nijs, M. Schick, and S. L. Keller, “Line tensions, correlation lengths, and critical exponents in lipid membranes near critical points,” Biophysical journal, vol. 95, no. 1, pp. 236–246, 2008.

